# Glaciers retreat, stickleback advance: variation in demographic histories of freshwater stickleback populations

**DOI:** 10.64898/2026.09.11.750921

**Authors:** Hanna Rosinger, Ryan Greenway, Cameron M. Hudson, David J. Janssen, Jakob Brodersen, Blake Matthews, Philine G. D. Feulner

## Abstract

When colonizing new habitats, the evolution of populations can be affected by their demographic history and structure, including their effective population sizes and the time and nature of their divergence from an ancestral population. The deglaciation of Southern Greenland following the last glacial maximum offers the opportunity to comparatively study colonization events of fish populations among lakes of varying size and altitude, the latter of which partially reflects their geological age and isolation. This study investigates the demographic history of threespine stickleback (*Gasterosteus aculeatus*) from 17 freshwater lake populations in comparison to a marine population sampled across four fjord sites in Southern Greenland using whole-genome data. Our findings suggest that divergence of freshwater from ocean populations was concurrent with ice sheet retreat and lake formation, which occurred ∼4-13 thousand years ago. In all 17 freshwater populations, demographic modeling revealed a population decline upon lake colonization and prolonged periods of gene flow between freshwater and marine populations. We also found that bigger lakes had larger effective population sizes. These findings emphasize the interplay between local landscape and demographic history in shaping the population genetic structure of Southern Greenland stickleback populations. Our results underscore the relevance of such small scale variation in demographic history across the landscape for understanding postglacial colonization into freshwater ecosystems.

**Tease text:** Inferred demographic histories of threespine stickleback populations from Southern Greenland elucidate how landscape geological history shaped colonization of freshwater lakes. Little variation in divergence times between marine and freshwater populations suggests lakes were colonised swiftly when the ice sheet retreated. Variation in freshwater effective population sizes can be explained by lake area, and gene flow between ocean and lakes was pervasive during freshwater population establishment.

## Introduction

Demographic variation has the potential to considerably affect evolutionary processes, by affecting genetic diversity and adaptation and the long-term survival probability of populations (Frankham, 1996; Hewitt, 2000; Lande, 1993; Slatkin 1987). Population bottlenecks and expansions can shape the genetic variation present within populations and influence subsequent evolutionary responses to environmental change (Ellegren and Galtier, 2016). For example, as observed in harvested populations of Northern elephant seals (Hoelzel et al., 2002). Past demographic processes such as changes in population size, isolation, and gene flow leave a unique molecular signature that permits the inference of such historical events and past evolutionary processes (Gutenkunst et al., 2009; Peter and Slatkin; 2013, Ray and Excoffier, 2009).

Inference about demographic processes entails estimating key population genetic metrics, such as effective population size (N_e_) and migration rate. N_e_ was defined by Wright (1931) for a single isolated idealised population in which generations do not overlap, population size is constant, mating is random and no selection occurs. The N_e_ of an actual observed population is the number of individuals in such an idealised population that experience the same extent of genetic drift as in the observed population (Futuyma, 2017). Hence, N_e_ determines the rate of loss of genetic diversity, the rate increase of inbreeding and the relative effectiveness of selection (Charlesworth et. al., 2009; Wang, 2005; Waples, 2002). Further, in the short term, changes in allele frequencies within a population depends on N_e_, except when migration is high, migration rate > 10% (Waples, 2025). However, less than one migrant per generation can already significantly boost the genetic diversity of a population, relative to a population without migrants (Waples, 2025). Complex demographic histories in natural populations that include migration and substructured populations make accurate estimates of contemporary N_e_ more difficult (Gilbert and Whitlock, 2015; Ryman et al., 2019). Nevertheless, N_e_ is important for understanding long-term population viability, as small effective population sizes are often associated with reduced adaptive potential and increased extinction risk (Frankham et al., 2014; James et al., 2016). Thus, the reconstruction of demographic histories sheds light not only on the role of selection in the past but may also enable us to predict future evolutionary trajectories of populations (Hohenlohe et al., 2021; Urban et al., 2016).

The phenotypic evolution of freshwater populations of threespine stickleback (*Gasterosteus aculeatus)* following their colonization from the ocean is a classic model of rapid adaptation, and occurs against a backdrop of complex demographic processes, potentially including population bottlenecks, changes in gene flow, and isolation events. The threespine stickleback is a prominent model organism in ecology and evolution (e.g., Bell and Foster 1994; Fang et al., 2018; Gibson, 2005) and has a wide range of tools and data available for discerning demographic variation. The species originated from the eastern Pacific and have dispersed through a trans-Pacific migration to the Atlantic (Artamonova et al., 2022; Fang et al., 2018). Although it is an ancestrally marine fish, it has adapted to freshwaters throughout the Northern hemisphere on multiple independent occasions since the last glacial maximum (Fang et al., 2018; Magalhaes et al., 2016; Mäkinen and Merila, 2006), often with exceptional speed and predictability (Barrett et al., 2011; Lescak et al., 2015; Roberts Kingman et al., 2021). Broad patterns of postglacial colonization of freshwater by sticklebacks are well documented in Northern Europe and North America (Makhrov et al., 2025), but we have a poor understanding about how the interplay between ecological heterogeneity of lakes (e.g., lake age, size, connectivity) and demographic processes influence the rate and mechanisms of population divergence. Southern Greenland offers an ideal location to study the demographic history of freshwater stickleback populations as the region was either fully covered by ice or submerged in the ocean during the last glacial maximum (∼22 thousand years ago (kya)), with lake formation in the early Holocene driven by retreat of the Greenland Ice Sheet and relative sea level changes. Lakes above ∼ 40 m were formed directly during deglaciation (ca. 10-12.5 kya, Leger et al., 2024; Levy et al., 2020) because their altitude was already above relative sea level of the time (Luetzenburg et al., 2026; Sparrenbom et al., 2006; 2013). Lakes below this level (∼ 40 m ± 10 m above current sea level) are slightly younger (8-11 kya), and formed when decreases in relative sea level concurrent with and since deglaciation, isolated them from marine environments (Luetzenburg et al., 2026; Sparrenbom et al., 2006; 2013). Marine threespine sticklebacks colonized those newly formed lakes and established freshwater populations. While some colonization events in Southern Greenland may have been marked by abrupt isolation, others might have occurred gradually, with varying connectedness and potential gene flow over time, and where gene flow might have occurred during divergence and isolation or been established later as a secondary contact. The potential for such variation in the landscape provides a useful model system to test how variation in demographic processes such as bottlenecks, divergence times, and gene flow shape contemporary populations. By focusing on the timing and mode of divergence, we aim to gain greater insights into the processes that govern colonization and establishment.

Currently, the only reconstructions of stickleback colonization in Greenland are from West Greenland populations, in which Liu et al., (2016; 2018) date the migration of sticklebacks to freshwater habitats and the associated population decline to approximately 3-9 kya. These analyses primarily reconstructed historical effective population sizes but did not estimate gene flow dynamics post-colonization. Similar patterns are observed in Greenlandic and other Northern European populations, where colonization of freshwater habitats is typically associated with a population decline followed by stabilization at a smaller population size (Liu et al., 2018).

Here, we present demographic modelling of 17 natural freshwater stickleback populations in Southern Greenland, and specifically make inferences about three demographic parameters: (i) the timing of divergence between marine and freshwater populations, (ii) changes in historical population sizes, and (iii) the timing and extent of gene flow between marine and freshwater populations. We produced 124 whole-genome-sequences (WGS) from 21 populations (17 freshwater and 4 marine sites) in Southern Greenland. We used *fastsimcoal2* (Excoffier et al., 2021) to investigate gene flow modes and population size changes in a two-population model (i.e., marine and freshwater) treating each lake as a unique colonization event. We tested multiple scenarios and our best fitting model includes both gene flow and allows for a change in the associated parameter estimates, allowing the model to infer scenarios of divergence with gene flow (continuous or ancient) as well as secondary contact after divergence.

## Methods

### Sampling and Sequencing

In total, samples from 124 individuals from 21 sites were collected in Southern Greenland in 2016 and 2019 (Collection Permits: G16-040 and G19-002, Ministry of Fisheries, Hunting and Agriculture, Government of Greenland; Table S1, S2; Figure 1 (A)). Fish were caught with minnow traps or gill nets and euthanized with MS-222. Fin clips were stored in ethanol for processing in Switzerland. DNA extractions were carried out using the DNeasy Blood & Tissue Kits. Illumina TruSeq PCR free libraries were prepared by the NGS facility Bern and paired-end (2× 150bp) sequenced in two batches, one batch on two lanes and the second batch on three lanes on an Illumina NovaSeq 6000. The raw data used for this study can be found under the bioproject PRJNA1240318.

**Figure 1.**
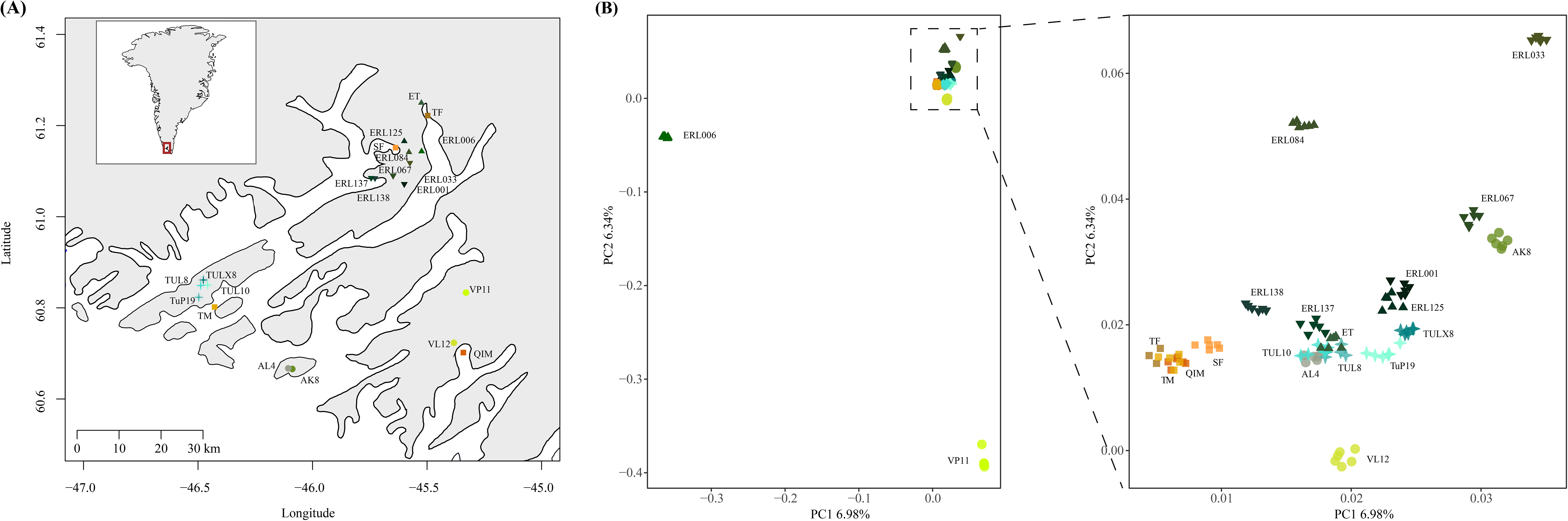
(A) Map of sampling locations and (B) PCA indicating clustering of individuals per population and differentiation among populations. Individuals are coloured according to their sampling site. Freshwater populations in circles, triangles, inverted triangles and stars grouped per geographic origin; marine populations in squares. The Sermilik Fjord samples were from a coastal lagoon site with resident stream and lake populations upstream of the site. Geographic locations and population abbreviations in Table S1.

### Read processing and genotype calling

After quality checking with fastqc (https://www.bioinformatics.babraham.ac.uk/projects/fastqc/) we removed adapters, retained reads with a quality phred scale >10, and trimmed poly-G tails. Reads were aligned to the latest stickleback reference genome (v.5, https://stickleback.genetics.uga.edu) with bwa (v.0.7.17; Li, 2013). We marked and removed duplicates with picard (v. 2.25.7). Additionally, we removed the sex chromosomes Y and XIX (the X-chromosome in the stickleback reference) as well as the mitochondrial genome and unknown chromosome sequences. As we detected a known inversion on chromosome XXI in our dataset, we excluded the entire chromosome because inversions affect recombination and alter drift (see Berdan et al., 2023). With this exclusion of regions we followed common best practice (Foote et al., 2016; Li & Durbin, 2011; Nadachowska-Brzyska et al., 2016). We excluded masked regions and used bcftools mpileup (v.1.12; Li, 2011) to call variants and monomorphic sites on reads with a minimum mapping quality of 20 and skipped indels in the process. The resulting vcf file contained 336,063,657 monomorphic sites and 8,030,216 SNPs (86.4% of genome). Single nuclear polymorphisms (SNPs) and monomorphic sites were filtered with bcftools and vcftools (Danecek et al., 2011) for a quality of 20, an individual depth of 4x, a depth filter per site across all individuals of the median ± sd (1631.97 ± 285.33) and a max missing filter of 0.5. Then we further filtered SNPs to keep only biallelic sites and applied a thinning threshold of 10bp. The filtered dataset contained 4,723,738 SNPs and 161,150,045 invariant sites (∼ 41.7% of the genome, Table S3). The average depth per sample was 13.02 (SD=3.76; Table S2).To construct site frequency spectra for demographic modelling we only used sites without any missing data in a population or a population pair.

### Population statistics

To assess population structure we calculated pairwise F_ST_ and nucleotide diversity π, using pixy (Korunes & Samuk 2021) in sliding windows of 10,000 bp. Further we utilised the R package SNPRelate (Zheng et al., 2012) to perform a principal component analysis (PCA) using a LD pruned set of variable sites (plink settings ––indep-pairwise 50 5 0.2; Purcell et al., 2007). All output data was processed and plotted with R (v.4.4.2; R Core Team; 2021).

### Demographic modelling and association with lake properties

We reconstructed effective population size trajectories (stairway plots) for each population with a method that has a resolution between about 40-50 kya (Nadachowska-Brzyska et al., 2022). We calculated folded site frequency spectra (sfs) with easySFS (https://github.com/isaacovercast/easySFS) and used the program stairway plot 2 (Liu and Fu, 2020) with the default settings for folded sfs using the following parameters: nseq: 12; L: sum of all sites – monomorphic and SNPs – without missing data, SFS: the sfs calculated for each population but excluding monomorphic sites (first entry of sfs), mutation rate: 3.7×10^−8^ or 5.11×10^−9^, year_per_generation: 1.

We used *fastsimcoal2* v.28. (Excoffier et al., 2021), a coalescent simulation based method, to explore seven demographic scenarios (Figure S2), using 2d folded sfs (calculated with easySFS) for pairwise combinations of all 17 freshwater with one representative marine population. All freshwater populations have a N=6, and in the parameter space we evaluate simulations suggest that accurate parameter estimates and model selection should be possible (Robinson et al. 2014). We selected the marine population QIM for modelling because the sampling effort was balanced with the freshwater populations (N=6), it was the most distant geographical location from all freshwater sampling sites, and the F_ST_ values between QIM and the freshwater populations were consistently high in comparison to the other marine sampling sites. In addition, we tested for a subset of selected freshwater populations how stable our parameter estimates are to this choice. For five freshwater populations we repeated the analysis using a mixed marine sample (two individuals of three marine populations). On the basis of reconstructions of lake formation ages for Southern Greenland (Luetzenburg et al., 2026; Leger et al., 2024; Levy et al., 2020; Sparrenboom et al., 2006; 2013;) and subfossil remains of sticklebacks found in Greenland (Bennike, 1997) we set the upper boundary of our models at 20 kya.We modelled seven scenarios (Figure S1, S2) estimating 4-9 parameters per scenario: a simple model with divergence without any gene flow and no changes in population sizes, three different gene flow models (continuous, ancient, or secondary contact), and two models that allow for changes in population size in the freshwater population after formation. The final model which we used allowed a change in migration rate after divergence. Introducing this potential change of migration rate and setting the parameter space to allow for the absence of gene flow allowed the model to estimate parameters that fit a scenario of divergence with ancient gene flow or a secondary contact scenario.

All input data (.tpl, .est, sfs, code) are provided in our data package (DOI: 10.25678/000E7B). Each demographic scenario was modeled 100 times. A global maximum likelihood estimate was calculated across those runs and the Akaike’s Information Criterion (AIC) was calculated for the best model for comparison. Additionally, we performed a maximum likelihood-bootstrapping with 1 million replicates for each model in *fastsimcoal2* and plotted the results in violin plots (Figure S3). Diversity estimate bootstrapping was done for the final model (model g) through non-parametric bootstrapping in *fastsimcoal2* by computing 100 bootstrapped sfs replicates. For each replicate, *fastsimcoal2* was run three times and the best run was chosen for the confidence interval calculations (Figure S4). Additionally, we plotted the sfs fit of the model in Figure S5.

To test if abiotic and physical lake properties explain variation in model-based demographic parameter estimates of effective population sizes for the freshwater populations (N_e_), divergence times (T_DIV_), across lakes we used linear regression (R command lm; Chambers, 1992; Wilkison et al., 1973). We assessed predictor pairs for collinearity (area and altitude, R command cor.test and car::vif) and right-skewedness, corrected for multiple testing with the Benjamini-Hochberg false discovery rate.

## Results

### Population statistics

We generated whole genome sequences for 124 individuals from 21 sampling locations in Southern Greenland (17 freshwater lakes with 6 individuals per lake and 4 marine sites with 3-7 individuals per site). Population diversity analyses revealed that sticklebacks in freshwater lakes showed only a minor reduction in population diversity (π = 0.0013-0.0033) in comparison to marine sites (π = 0.0035-0.0036) (Table S4). Pairwise F_ST_ between freshwater lakes was higher (F_ST_ = 0.067-0.594) than between marine sites (F_ST_ = 0-0.0237; Table S5). A PCA (Figure 1 (B)) showed that individuals clustered well by lake, although the first two PC axes explain only a small part of the overall variation (PC1 6.98 %, PC2 6.34 %). Principal component 1 separated most freshwater lakes from the marine population. Two populations (VP11 and ERL006) stand out as the most distinct clusters, likely indicating a pronounced influence of genetic drift in those two populations. While most lake populations differentiate from the marine on PC1, ERL84 overlaps with the marine on PC1 but separates well on PC2.

### Demographic modelling

We used stairway plots to infer trajectories of effective population size (N_e_) for all 21 populations. Based on our field observations, a generation time of 1 year is most plausible for freshwater sticklebacks in Greenland, meaning that the “generations before present” is equivalent to the “number of years ago”. While we modeled all 21 populations independently, their trajectories converge to similar N_e_ at about 50 thousand years ago (kya) ranging between 20-50 thousand individuals (Figure 2), as expected for populations that shared a common ancestor. Ten of the 17 freshwater populations show a single decline either steeply (six lakes) or more steadily (four lakes) just after ∼5 kya (Figure 2 (B)). In the other seven, population sizes remain rather constant around that time with a more gradual reduction in N_e_ occurring as recently as 1-2 kya (Figure 2 (C)). Of those seven, four populations show an initial reduction around 30 kya additionally, well before the current lakes were formed, and then a recovery of N_e_, around 5-10 kya and prior to declining characteristics for all freshwater populations. All marine populations maintain a high N_e_, which increased slightly around 5-10 kya (Figure 2 (D)), around the time when the ice sheet retreated in Southern Greenland (Leger et al., 2024; Levy et al., 2020) and also show a slight initial reduction around ∼30 kya.

**Figure 2.**
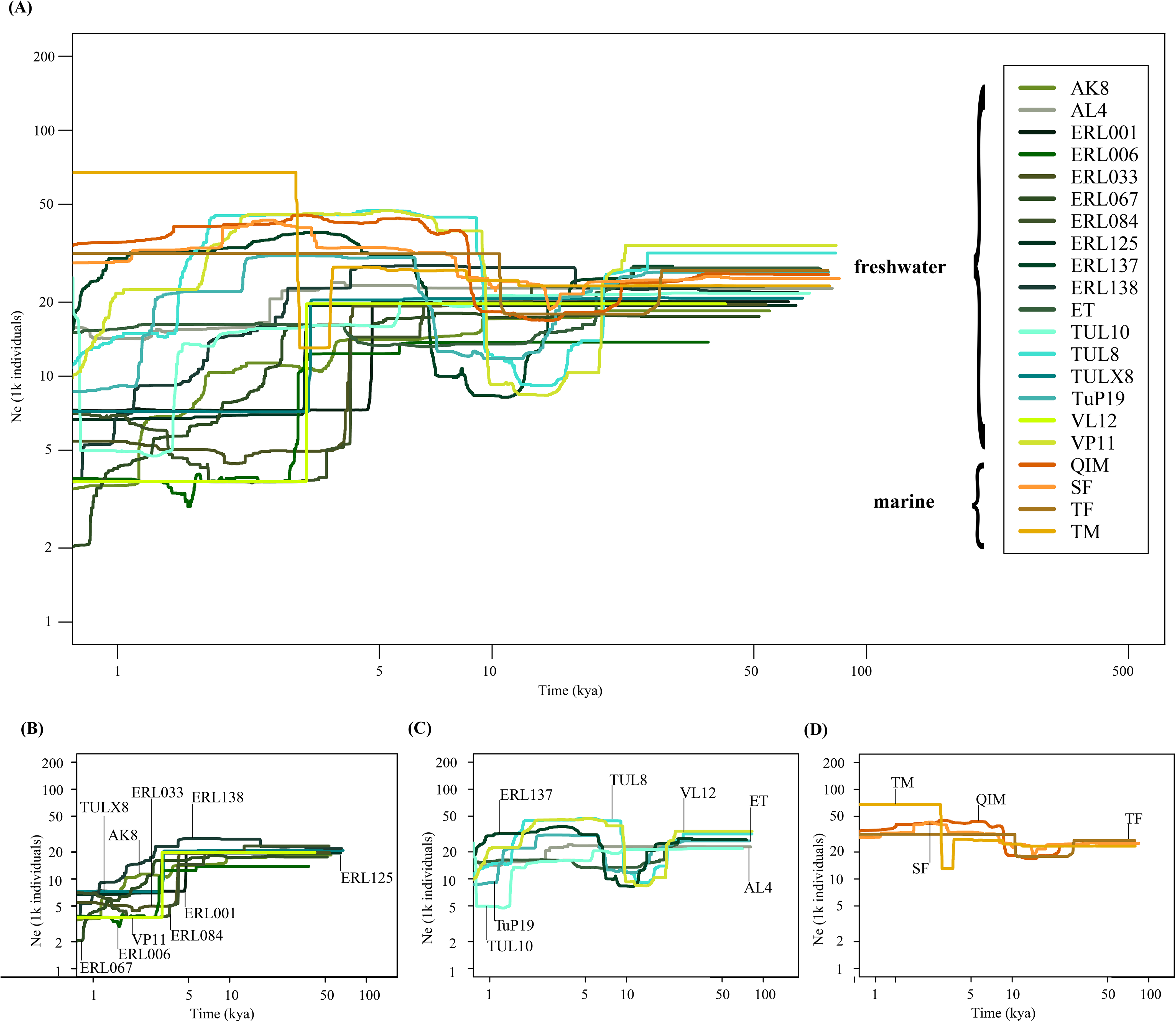
Trajectories of effective population sizes (N_e_) among populations. Stairway plots using a mutation rate of 3.7×10^−8^. Time in ky is shown on the x-axis, with effective population size on the y-axis. Panel (A) shows all populations analysed, (B) freshwater populations characterised by a single decline ∼5 kya (C) freshwater populations a rather constant Ne ∼5kya and a decline ∼1-2 kya, and (D) four marine population trajectories that match the expectations of a stable marine population over time. Individual plots of each population in Supporting Information Figure S6. Geographic locations and population abbreviations in Table S1.

To more explicitly test demographic scenarios and estimate parameters including a time of divergence between freshwater and marine populations we made use of a model-based approach as implemented in *fastsimcoal2*, testing multiple scenarios (Figure S2, Table 1, S6, S7, different mutation rate Table S8, different outgroup Table S9). Across the 17 lakes the simple model with no gene flow (model a), and the two models that allowed the freshwater population to change size after divergence (models e and f) as well as the model with continuous gene flow after divergence (model b) were less likely than at least two of the other gene flow models (models c, d, and g) (Figure S3). To allow a comparison of parameter estimates across lakes we used model g (Table 1, Figure S4), which combines the two alternatives of divergence with ancient gene flow and a secondary contact. This was achieved by setting the parameter space to allow for a change in gene flow. The sfs fit can be found in Figure S5.

**Table 1.** Demographic parameter estimates vary among populations. Results of the bootstrapping of the final model (model g) of *fastsimcoal2* with a mutation rate of 3.7× 10^−8^ and K = 9, which is the number of parameters in the *fastsimcoal2* model. All parameters are mean values of the bootstrapping. Effective population sizes (N_e_) of ancestral, freshwater, marine, N_1_M_21rec_= effective migrants from marine to freshwater after CHANGEM (present), N_1_M_21old_= effective migrants from marine to freshwater before CHANGEM (past), N_2_M_12rec_= effective migrants from freshwater to marine after CHANGEM (present), N_2_M_12old_= effective migrants from freshwater to marine before CHANGEM (past), T_DIV_= divergence time in kya, CHANGEM= timing of migration matrix change in kya. Extended results can be found in Supplementary information Table S6. Geographic locations and population abbreviations can be found in Table S1.

| Population | $N_e$ ancestral | $N_e$ freshwater | $N_e$ marine | $N_1M_{21rec}$ | $N_2M_{12rec}$ | $N_1M_{21old}$ | $N_2M_{12old}$ | $T_{DIV}$ | CHANGEM |
| --- | --- | --- | --- | --- | --- | --- | --- | --- | --- |
| AK8 | 47519 | 2676 | 56165 | 0.19 | 2.66 | 1.50 | 0.17 | 7.06 | 0.86 |
| AL4 | 47625 | 11109 | 49009 | 3.41 | 2.58 | 0.11 | 0.49 | 8.63 | 3.14 |
| ERL001 | 47398 | 3690 | 56314 | 0.51 | 3.30 | 1.73 | 0.26 | 7.46 | 1.31 |
| ERL006 | 47133 | 1192 | 64188 | 0.12 | 0.30 | 5.91 | 0.65 | 6.12 | 1.22 |
| ERL125 | 46877 | 3272 | 52355 | 0.65 | 1.64 | 0.51 | 0.25 | 12.65 | 4.42 |
| ERL137 | 47489 | 5311 | 53385 | 1.71 | 1.32 | 0.20 | 0.26 | 8.42 | 3.56 |
| ERL138 | 48112 | 3680 | 64089 | 0.44 | 0.09 | 5.39 | 0.30 | 3.61 | 0.68 |
| ERL033 | 47929 | 2300 | 57618 | 0.06 | 0.71 | 1.69 | 0.08 | 5.57 | 0.83 |
| ERL067 | 47650 | 2110 | 57710 | 0.19 | 0.86 | 1.60 | 0.10 | 6.60 | 0.85 |
| ERL084 | 47411 | 2126 | 62473 | 0.28 | 0.86 | 3.75 | 0.23 | 5.85 | 1.35 |
| ET | 47614 | 4629 | 54761 | 1.18 | 1.53 | 0.69 | 0.44 | 7.92 | 3.56 |
| TUL10 | 46962 | 6756 | 49530 | 1.52 | 2.04 | 0.08 | 0.08 | 10.53 | 4.31 |
| TUL8 | 47647 | 10147 | 51835 | 3.61 | 1.80 | 0.22 | 0.47 | 7.19 | 2.92 |
| TULX8 | 47582 | 3777 | 53276 | 0.66 | 2.25 | 1.05 | 0.41 | 10.49 | 2.84 |
| TuP19 | 47714 | 6674 | 51086 | 2.09 | 1.36 | 0.16 | 0.21 | 8.90 | 4.14 |
| VL12 | 47575 | 11372 | 48527 | 3.44 | 2.54 | 0.13 | 0.38 | 9.24 | 2.90 |
| VP11 | 47673 | 1084 | 60685 | 0.04 | 0.10 | 3.49 | 0.16 | 6.58 | 0.97 |

The results of this final model demonstrate that all freshwater populations have experienced a population size reduction after divergence from the marine population. The same marine population (QIM) was used across all simulations with different freshwater populations, and consequently the marine and ancestral population sizes are similar across simulations (Figure 3 (B)). This also fits with the stairway plots where the marine populations are stable across time (Figure 2 (D)). The freshwater population sizes are smaller than the marine across all populations, but estimates varied across lakes between 1084 (VP11) and 11372 (VL12) (Table 1). The timing of the divergence from marine to freshwater in the *fastsimcoal2* modelling was dated to 4-13 kya (mutation rate 3.7×10^−8^, Table 1, Figure S4), a dating which is well bounded by regional lake formation ages. Interestingly, time of divergence correlated only marginally and not significantly with the N_e_ of the freshwater population (Figure 3 (A), p=0.059), hence other factors than the colonization age seem to determine the freshwater population size. Using a mix of marine samples instead of QIM confirmed most parameter estimates only time of divergence decreased slightly across all populations (Table S9).

**Figure 3.**
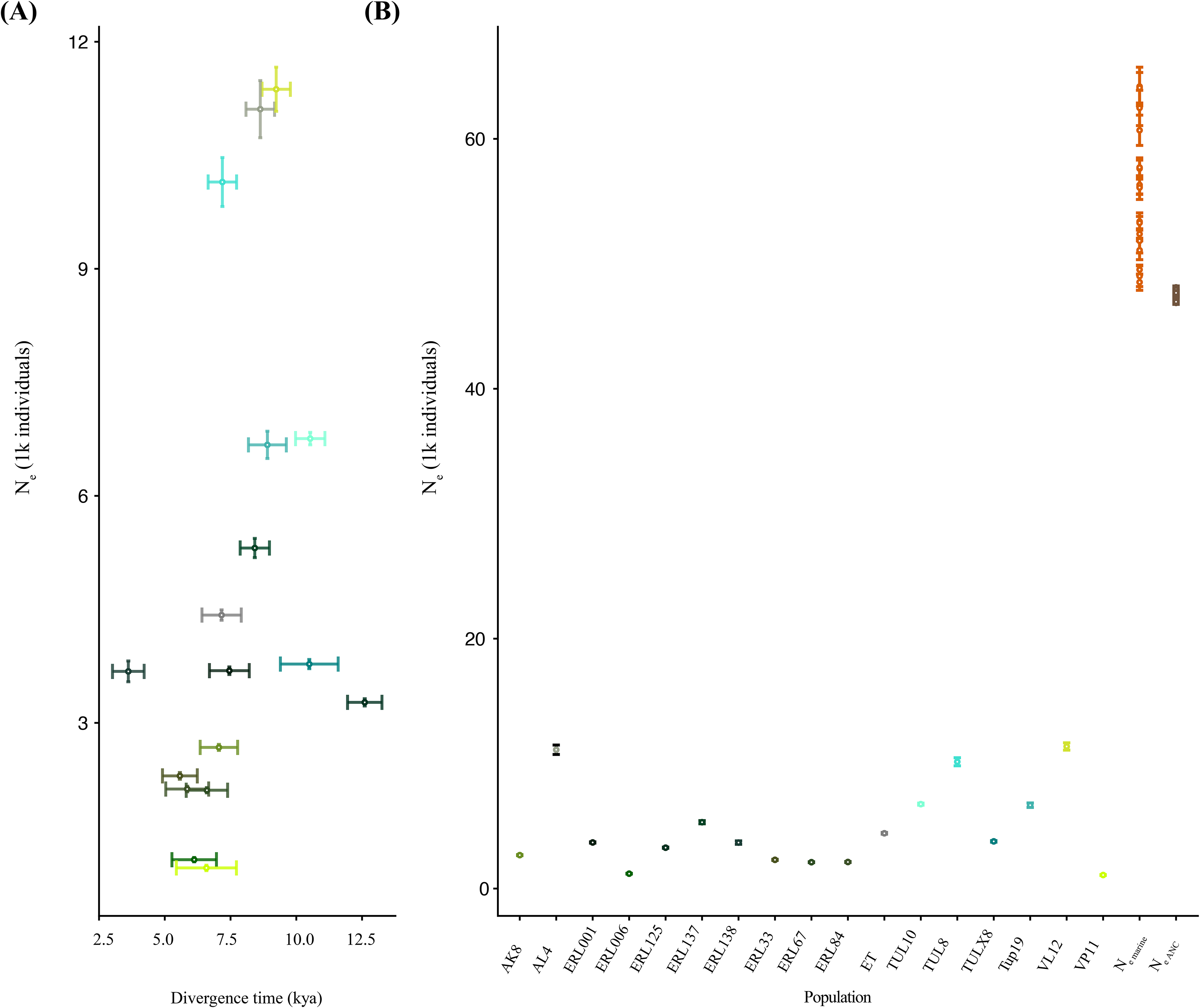
Variation in effective population sizes and time of divergence estimates among populations. (A) Divergence time (T_DIV_) of each freshwater population plotted against the effective population size (N_e_ freshwater) show no correlation (β=536.13, SE=370.31, adjusted p=0.168), average and 95 % confidence interval are presented for both T_DIV_ and N_e_. (B) Average and 95% confidence intervals are presented for effective population sizes of either freshwater populations, marine population or the ancestral population. Geographic locations and population abbreviations in Table S1.

Our results further showed that while gene flow improved model fit (Figure S3), estimates were rather low, however lakes differed in the respect of when gene flow occurred relative to divergence. Eight lakes had a lower migration rate from marine to freshwater right after divergence and migration increased more recently suggesting a secondary contact of these freshwater populations with the ocean (Figure 4 (A), S7), and in nine lakes the pattern was reversed suggesting past gene flow right after divergence (Figure 4 (B), S7). Only for lakes ERL125 and TULX8 confidence intervals overlap; hence, those populations showed no significant change in migration. For the eight secondary contact lakes, migration from freshwater to marine matched the pattern detected in the other direction (Figure 4 (C), S8). Lakes with higher past marine gene flow, did not reflect that pattern in the freshwater-to-marine migration; most lakes (6 of 9) showed a low-to-high migration pattern, while three had overlapping confidence intervals (Figure 4 (D), S8).

**Figure 4.**
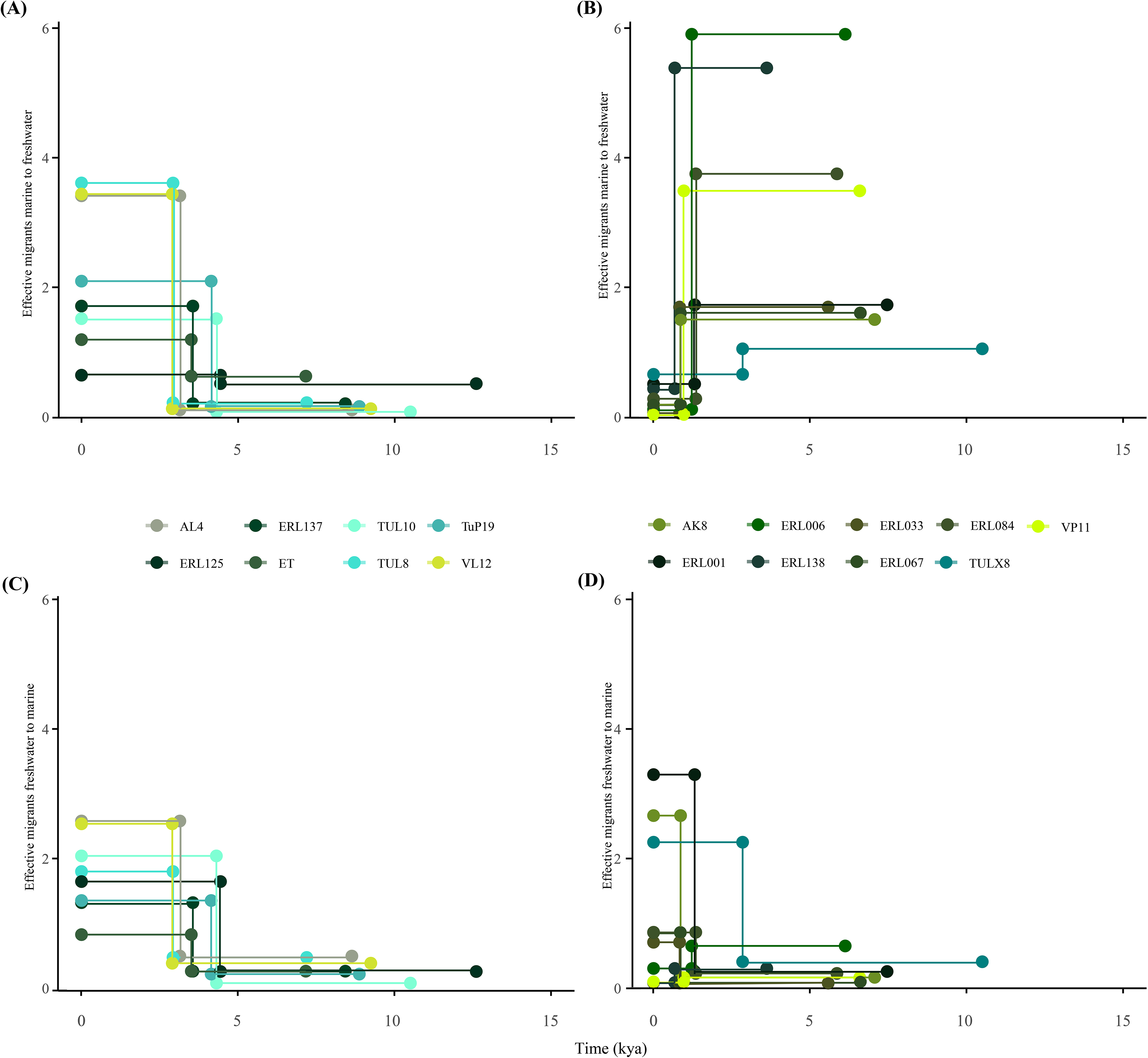
Variation in effective migrants among populations. (A) and (B) effective migrants (per generation) from marine to freshwater, categorized due to the direction of change as secondary contact (A) or divergence with gene flow patterns (B). (C) and (D) effective migrants from freshwater to marine. Geographic locations and population abbreviations in Table S1.

### Association of lake properties and demographic parameter estimates

We tested if lake altitude and area explain some variation in demographic parameter estimates. Altitude was not a good predictor of divergence time (Figure 5 (A), β=−8.76, SE=8.45, adjusted p=0.352). However, altitude explains variation in N_e_ and migration well (Figure 5 (C), β=−21.22, SE=6.63, adjusted p=0.013), with higher altitude lakes being more distant from, and more poorly connected to, the marine population and therefore less genetically diverse (lower N_e_). Lake area explained some variation in both demographic parameters, with the larger lakes being slightly older, and having a larger effective population size (Figure 5 (B), β=1004.51, SE=1042.93, adjusted p=0.352; (D), β=4171.66, SE=817.62, adjusted p<0.001). In our study lakes, altitude and area are not correlated, and so these lake features might have independent effects on demography (VIF=1.09; Pearson’s correlation r=0.26, df=15, p=0.31).

**Figure 5.**
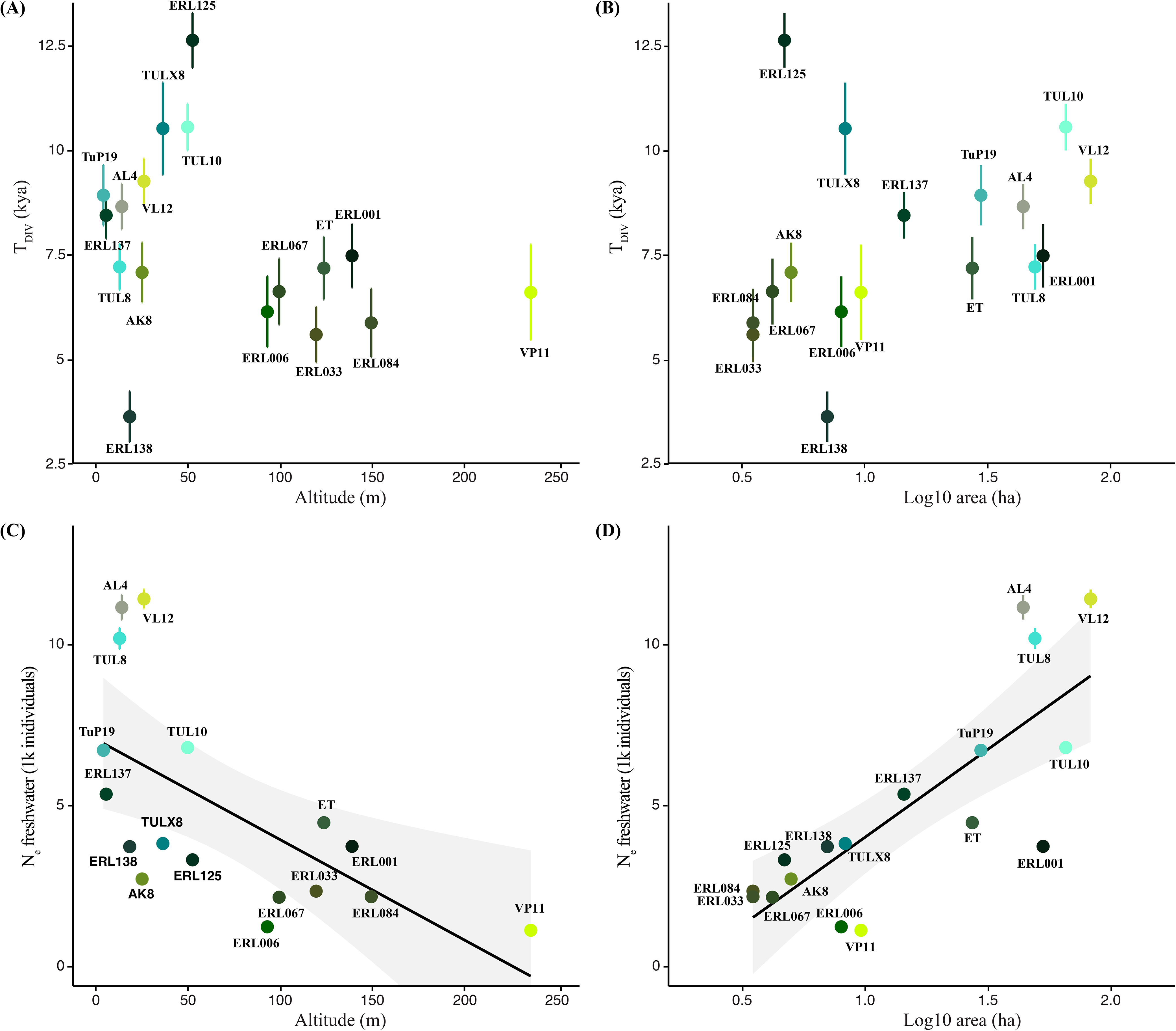
Variation in lake parameters (altitude, area) explain variation in demographic parameters, divergence time (T_DIV_) and effective population size of freshwater populations (N_e_ freshwater). (A) Regression of altitude (β=−8.76, SE=8.45, adjusted p=0.352) or (B) log10 area (β=1004.51, SE=1042.93, adjusted p=0.352) and divergence time, (C) regression of altitude (β=−21.22, SE=6.63, adjusted p=0.013) or (D) log10 area (β=4171.66, SE=817.62, adjusted p<0.001) and N_e_ freshwater. The plots show parameter estimates (bootstrap means and confidence intervals) and raw values of lake properties while statistics are based on residuals. Geographic locations and population abbreviations in Table S1.

## Discussion

The freshwater sticklebacks of Southern Greenland are an ideal system to study how local-scale variation in demographic histories is related to evolutionary responses of populations colonizing novel environments. The recent landscape history of Greenland is well studied at regional scales, for example in terms of ice sheet retreat (Leger et al., 2024; Levy et al., 2020) and relative sea level fall (Luetzenburg et al., 2026; Sparrenbom et al., 2006; 2013), both of which drive lake formation, and this provides a useful starting point for studying general patterns of population dynamics and demographic histories (Luetzenburg et al., 2026). However, at local scales, the specific positioning of lakes in the landscape and their ocean connectivity via migration, there is still considerable uncertainty about the timing of population establishment and the extent of gene flow during the ongoing process of stickleback colonization of freshwater environments.

### Variation in demographic histories among lake populations

Historical declines in N_e_ can be relevant for population persistence and adaptation to change and are thus important to recognize and study (Charlesworth 2009). While N_e_ trajectories were relatively stable for marine populations (Figure 2 (D)) freshwater populations showed more heterogeneity (Figure 2 (B), (C)). This is in line with past work, in which stable marine population sizes are documented (Liu et al., 2016; 2018) and declines of freshwater N_e_ in the last 10 ky, indicate colonization of freshwater habitats and cessation of gene flow with marine populations (Coll-Costa et al., 2024).

A decline of N_e_ all freshwater populations is also supported by the demographic model that best fits the data and the corresponding parameter estimates, namely smaller N_e_ for all freshwater populations than for the ancestral and marine population (Figure 3 (B)). Additionally to a more recent decline, four freshwater populations showed a notable dip in their N_e_ trajectory between 10-30 kya (Figure 2, S6); a dip less pronounced but also noticeable in three marine populations. This dip occurred during the late pleistocene, just prior to (or concurrent with) the last glacial maximum (16-24 kya; Funder et al., 2011). Although speculative, such patterns could indicate that the ancestral population became locally confined to a glacial refugium before freshwater lake colonisation marked the decline in N_e_ (i.e., ∼12 kya; Leger et al., 2024; Levy et al., 2020;).

Island colonization studies, analogous to lake colonization, commonly observe a reduction in N_e_ following a split from the ancestral population (evidential in our demographic modelling) (James et al., 2016; Losos and Ricklefs, 2009). For example, white-footed mice in the Boston Harbor archipelago experienced significant reductions in N_e_ following isolation due to rising sea levels after the Last Glacial Maximum (Howell et al., 2025), as did vole populations following the human introduction to Orkney 5 kya and successive colonizations of smaller islands (Wang et al., 2023). While a population decline reduces genetic diversity and adaptive potential by increasing genetic drift, thereby altering the trajectories of colonizing populations (Kimura, 1983; Wright, 1931), it can also have no lasting impacts on molecular evolution (James et al., 2016). This was seen in Orkney voles where soft selection, relaxed environmental pressure and rapid reproductive rate mitigated the negative effects of inbreeding and drift (Wang et al., 2023). This might raise the expectation that divergence time and N_e_ are positively correlated, which is not the case for freshwater stickleback populations in Greenland (Figure 3 (A)). The lack of correlation is not surprising, as lake formation occurred at similar times for high elevation lakes (> ∼40 m, during glacial retreat at ∼10-13 kya; Leger et al., 2024; Levy et al., 2020) and low elevation lakes, which were isolated during relative sea level fall (8-11 kya; Luetzenburg et al., 2026; Sparrenbom et al., 2006; 2013). Instead, the contemporary model derived N_e_ varies across lake populations and N_e_ is best explained by combination of lake area and altitude (R^2^ =0.75,F =25.11). Effective population size increased significantly with lake area (p<0.001, Figure 5(D)) and decreased significantly with altitude (p=0.006, Figure 5(C)). So, larger, low altitude lakes tend to sustain larger populations, which makes them less prone to genetic drift and more likely to retain adaptive genetic variation (Frankham, 1996). Whereas smaller populations in small high altitude lakes are more susceptible to drift and the loss of adaptive genetic variation (Lande, 1993). However, additional empirical work is needed to test these mechanisms directly. A comparable trend has been observed in birds colonizing islands, where genetic diversity was positively correlated with island size, and low genetic diversity could be explained by population decline (Brüniche-Olsen et al., 2019; Petren et al., 2005).

Divergence time estimates (4-13kya) are concordant with both fossil (9 kya; Bennike, 1997) and geological evidence (9-13 kya; Luetzenburg et al., 2026; Leger et al., 2024; Sparrenbom et al., 2006; 2013). Other studies modelling stickleback colonization in Greenland also suggested a colonization event between 3-11 kya (Liu et al., 2016; 2018). These estimates are similar to the ones presented here. However, we note that we used the same mutation rate as the studies on Greenland stickleback to which we compare (Liu et al., 2016, 3.7×10^−8^), and which resulted in a reasonable dating given previous geological inference about the timing of lake formation in the study area. In contrast, using a lower mutation rate of 5.11×10^−9^ (Zhang et al., 2025) resulted in divergence time estimates predating established geological records in Greenland (Figure S9 and S10, Table S8). Admittedly the choice of mutation rate is highly relevant for concordance with the geological evidence and hence the concordance is based on the choice and not independent evidence.

### Limited but relevant gene flow variation amongst lakes

All best-fitting models included some degree of gene flow, highlighting the importance of gene flow during freshwater colonization. Therefore, although freshwater lakes currently are spatially isolated populations in the landscape, their history allowed for gene flow with marine populations. Based on the gene flow estimates we identified two distinct patterns suggesting either a divergence with (some) gene flow or a secondary contact scenario. These scenarios are marked by nearly contemporaneous but opposing changes in marine to freshwater migration (∼2-4 kya, Figure 4 (A), (B)), consistent with environmental changes in South Greenland at this time – namely rising relative sea levels beginning ∼4-5 kya (Luetzenburg et al., 2026; Sparrebom et al., 2006; 2013) and cooler, drier climate, peaking ∼3 kya (Andresen et al., 2004; Massa et al., 2012; Larsen et al., 2016; Schneider et al., 2024). A cooler and drier climate likely restricted connectivity of lakes from marine systems through longer annual ice coverage and possibly decreased lake volume and outlet stream flow. Lakes showing a decrease in marine to freshwater migration (Figure 4 (B)) are smaller (≤ 8.6 ha) and mostly higher elevation (all except AK8 and ERL138 are above 90 m; see Table S1). The two lower elevation lakes are also quite shallow (maximum depths of 2 and 1 m for AK8 and ERL138). These factors increase susceptibility to negative impacts from a cooler and more arid climate. In contrast, most lakes showing an increase in marine to freshwater migration (Figure 4 (A)) are lower elevation (all but ET are ≤ 50 m elevation) and larger (≥ 13 ha, Table S1). A cooling climate likely had less impact on these lakes, where the ice cover season is already shorter and habitat volume is larger, while rising sea levels had a larger relative effect on connectivity. Put simply, a 5 m sea level rise removes a larger percentage of the travel path for a lake at 15 m elevation than one at 100 m. Therefore, increased connectivity appears to outweigh the negative influence of climate in these larger, lower elevation systems. These shifts in migration coincide with shifts in N_e_. All of the lakes with a reduced marine to freshwater migration (Figure 4 (B)) show a decrease in N_e_ at the same time as the change in migration shift occurred (Figure 2 (B)). In contrast, lakes showing an increase in marine to freshwater migration (Figure 4 (A)) show a more stable N_e_ around that time (Figures 2 (C); except ERL125). Overall, lakes with an increase in marine connectivity show a significantly larger freshwater N_e_ (mean = 8000 compared to mean =2368; Welch’s t-test: t=-5.02, p=0.001). This suggests that connectivity facilitated a consistent influx of alleles and with it a larger effective population size, which in turn shielded the population from the negative genetic consequences in the newly founded populations (Ingvarsson 2001; Lowe and Allendorf, 2010). To maintain similar allele frequencies in marine and freshwater populations (i.e., hinder divergence), migration would need to be higher (> 10 effective migrants per generation) than most parameter estimates here (Lowe & Allendorf, 2010). However, harmful local inbreeding might already be avoided by >1 effective migrant per generation and, over time, small amounts of migration (< 1 effective migrant per generation) increase heterozygosity and are sufficient to spread advantageous alleles (Lowe & Allendorf, 2010; Waples, 2025). So gene flow estimates for Southern Greenland populations might indeed be relevant for future trajectories of individual lakes. Gene flow in the opposite direction (freshwater to marine), though low (1-5 effective migrants per generation, Table 1), might also be sufficient for freshwater adaptive alleles to flow back to the marine population and enhance its adaptive potential for future freshwater colonization events. Such gene flow is a key assumption of the transporter hypothesis (Schluter and Conte, 2009), in which freshwater alleles are maintained in marine populations at a low frequency, through a lasting influx of freshwater migrants. This process fuels the standing genetic variation necessary for rapid and repeated freshwater adaptation of sticklebacks across the Northern hemisphere (Galloway et al., 2020; Kingman et al., 2021). However, estimates of rates of gene flow that support the transporter hypothesis have been lacking so far. Overall, these findings highlight the interplay between geological history and demographic processes and their relevance for the contemporary evolution of populations.

## Impact summary

Following the retreat of the glaciers in Southern Greenland, after the last glacial maximum, newly formed freshwater lakes were colonized by threespined sticklebacks. This created a unique opportunity to study evolution in multiple replicates. Here, we reconstructed the demographic past of multiple populations and show that all populations declined in N_e_ during freshwater colonization which coincided with the ice sheet retreat in Southern Greenland. Further, gene flow and geographic constraints (e.g., lake area) are primary drivers of variation in effective population sizes. Our findings provide an empirical link between landscape features and demographic histories of colonizing species.

## Author contribution

HSR analysed the data with the support of PGDF and BM. The fish were sampled by RG, CMH, JB, and BM. RG conducted DNA-extraction. HSR wrote the manuscript with the support of PGDF and BM and input from all authors.

## Acknowledgments

We thank Rebecca Oester, Daniel Steiner, Jukka Jokela, Marvin Moosmann, Danina Schmidt, Rebecca Best, Marek Svitok and Zixin Li for their assistance and support in the field. We acknowledge the excellent local support of Ellen and Karl Frederiksen, who facilitated multiple field campaigns. This study was computationally supported by Niklaus Zemp and the Genetic Diversity Centre Zurich and the sequencing data was generated at the Next Generation Sequencing Platform at the University of Bern. The authors also like to thank Vitor Sousa and David Marques for their helpful insights and suggestions regarding *fastsimcoal2*.

## Figure captions for Supporting Information

**Figure S1.**
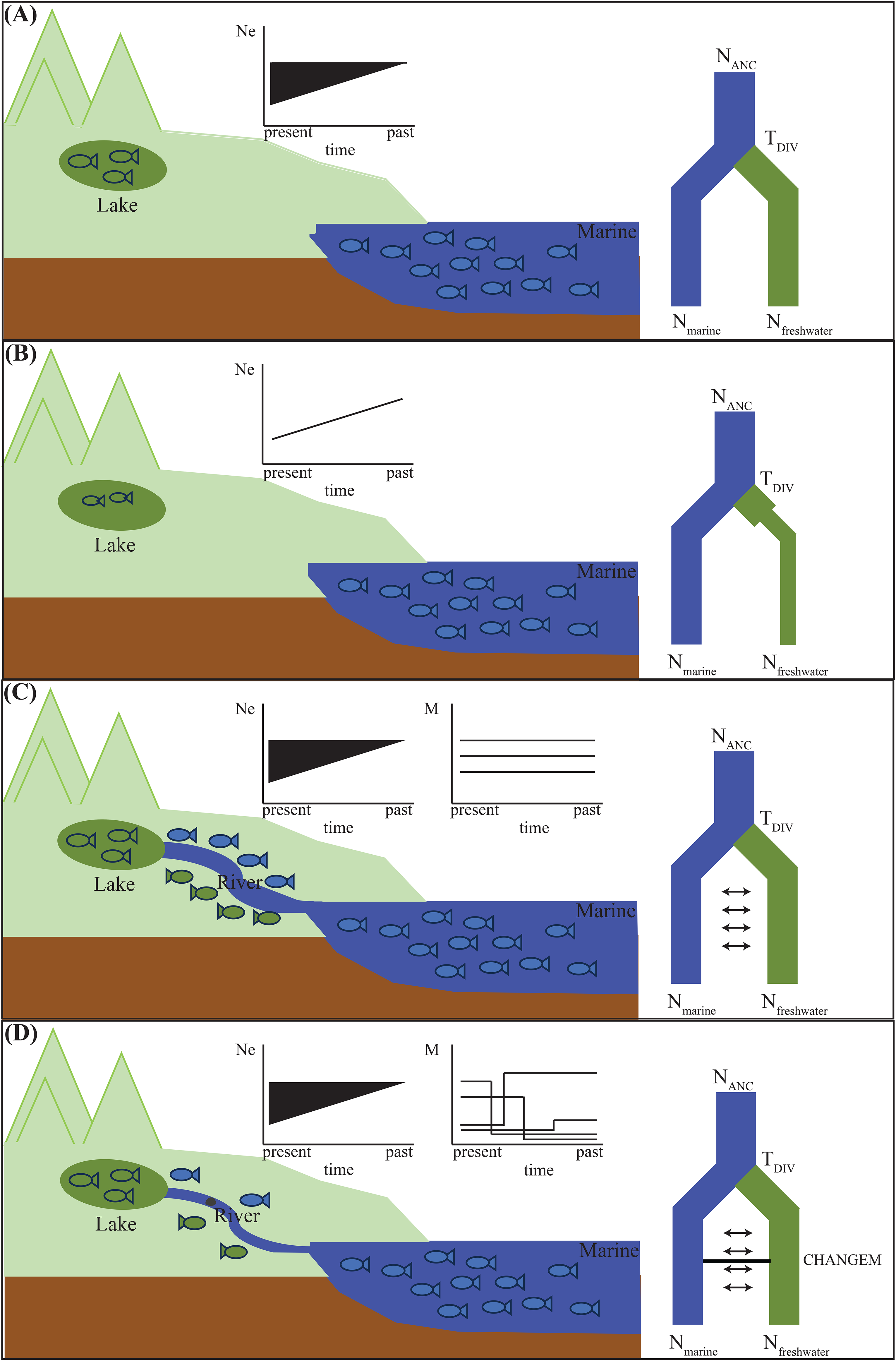
Conceptual figure of the demographic modelling and reasoning behind models which were tested in *fastsimcoal2*. The landscape is depicted with a green lake into which stickleback migrated and formed a freshwater population divergent from the marine population in blue. In (A) no connection of freshwater and marine population is present, the model is testing solely the time of divergence (T_DIV_) between the two populations and effective population (N_e_) decline is possible. In (B) we specifically model a decline in the freshwater population, and in (C) freshwater and marine populations are connected through a river which enables a constant gene flow between the populations, allowing for a decline in N_e_. In (D) freshwater and marine populations are connected through a river, the migration matrix in this model is allowed to change at one point (CHANGEM). In this case, the CHANGEM and settings are set so that the migration between the two populations can be very low; and a decline in N_e_ is possible

**Figure S2.**
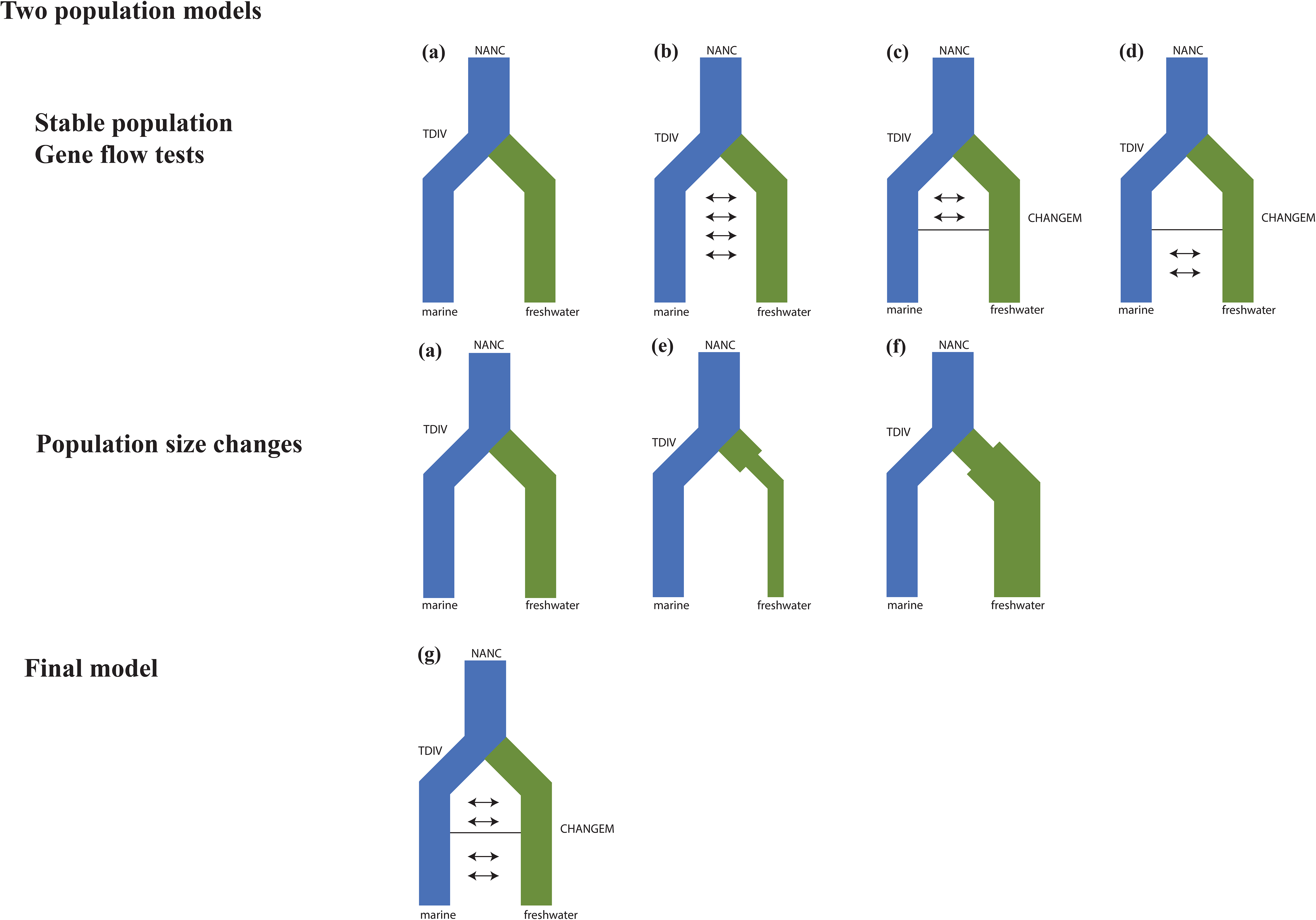
Models tested with *fastsimcoal2*. First row two populations gene flow models between marine and freshwater with stable population size, population size fluctuation possible but not forced. Second row population size changes of the freshwater population are enforced by the model. Third row-the final model combining gene flow and population size changes.

**Figure S3.**
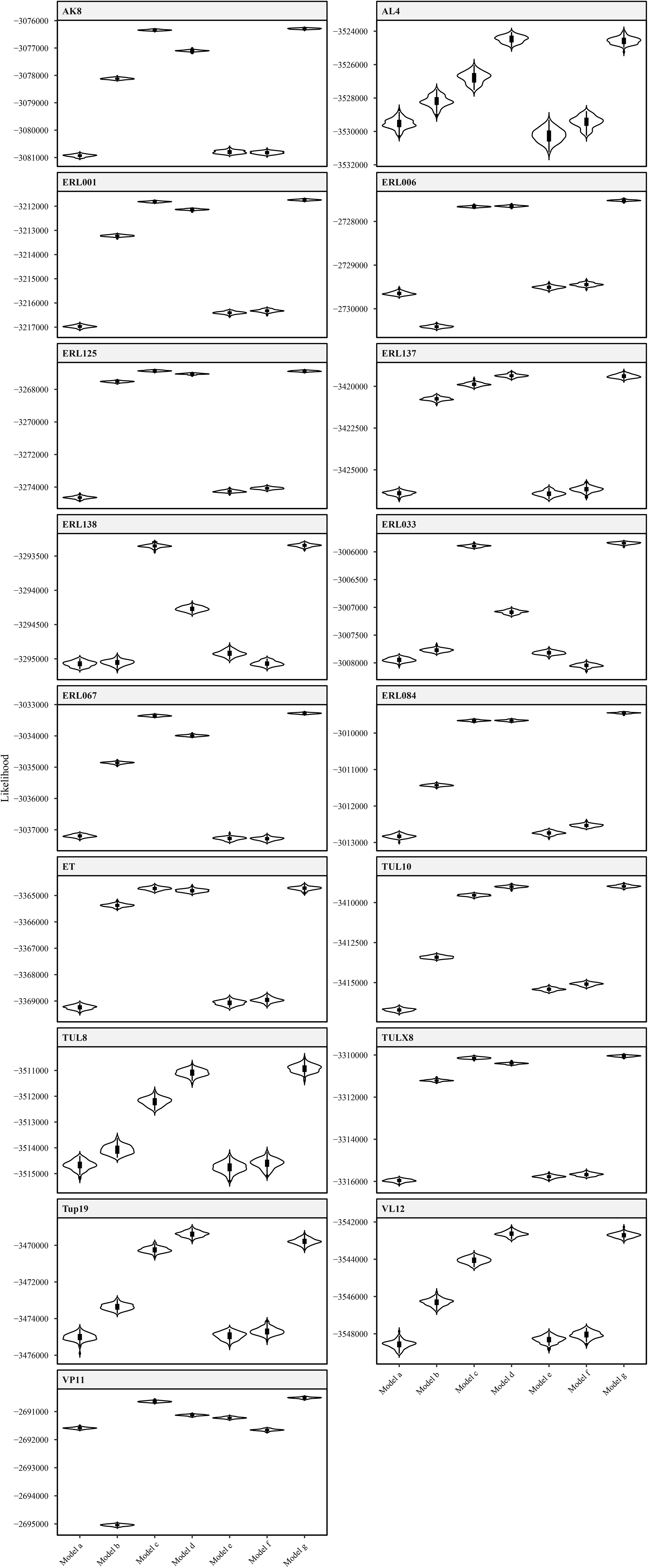
Likelihood distributions of all tested demographic models with the mutation rate 3.7×10^−8^. Violin plots show the distributions of likelihoods of 100 site-frequency spectra simulated under the maximum likelihood parameters inferred using the observed data. Note: An overlap of the likelihood distributions indicates an equal well fit to the observed data. Geographic locations and population abbreviations in Table S1.

**Figure S4.**
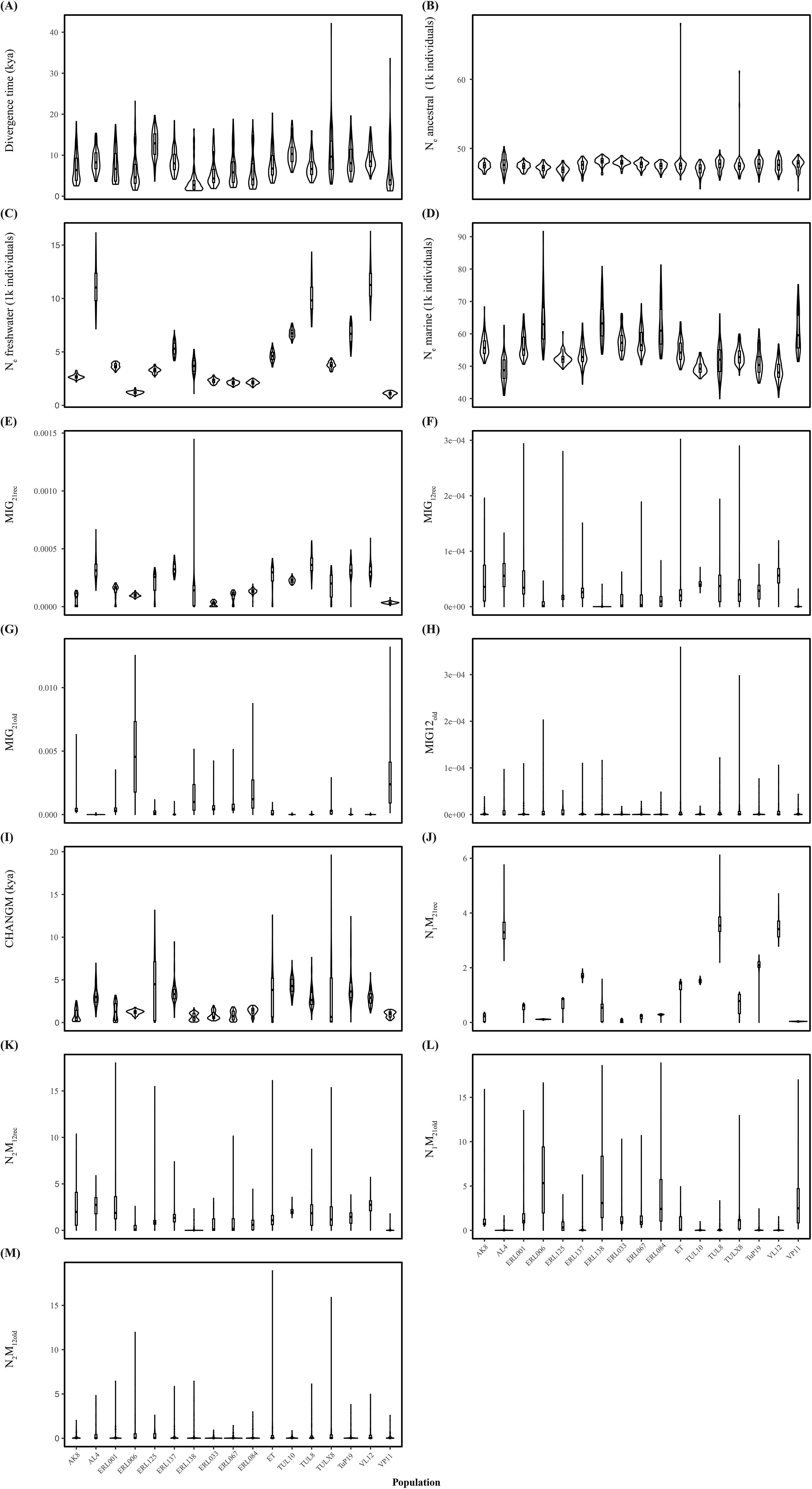
Violin plot of bootstrapped parameter estimates of the final model of *fastsimcoal2* (model g). (a) T_DIV_ between marine and freshwater population, (b) ancestral population size, (c) current N_e_ of the freshwater population, (d) current N_e_ of the marine population, (e) past migration from freshwater to marine, (f) past migration from marine to freshwater, (g) recent migration freshwater to marine, (h) recent migration from marine to freshwater, (i) time of migration change, (j) number of individuals migrating from freshwater to marine before change of migration, (k) number of individuals migrating from marine to freshwater before change of migration, (l) number of individuals migrating from freshwater to marine after migration change, (m) number of individuals migrating from marine to freshwater after change of migration. Violin plots depict parameter estimates of bootstrap replicates, with 95 % confidence intervals in green and blue dots representing the point estimates of the best model. Geographic locations and population abbreviations in Table S1.

**Figure S5.**
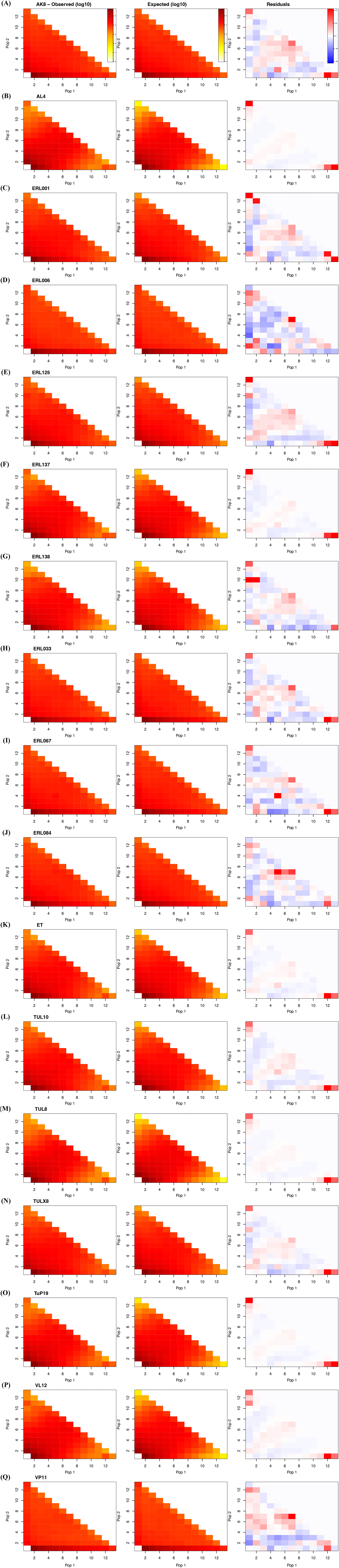
Observed and expected two-dimensional side frequency spectra for each population combination run of *fastsimcoal2* as well as the residuals fitting for the expected spectrum to the observed one. Population abbreviations in Table S1.

**Figure S6.**
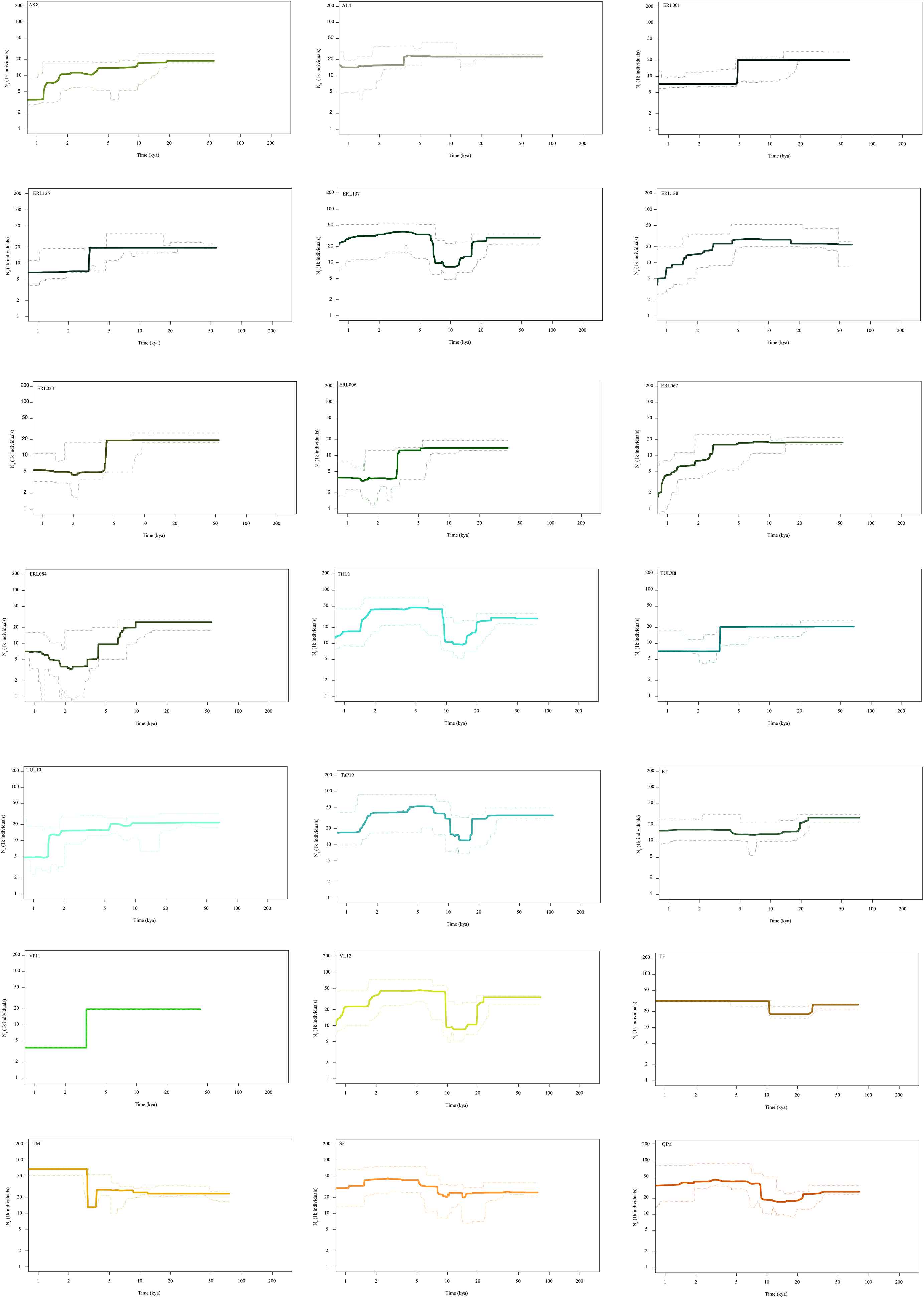
Stairway plots of effective population size (N_e_) for each population. The bold line represents the mean and the dotted line the 95% confidence intervals. A mutation rate of 3.7×10^−8^ was used. Geographic locations and population abbreviations in Table S1.

**Figure S7.**
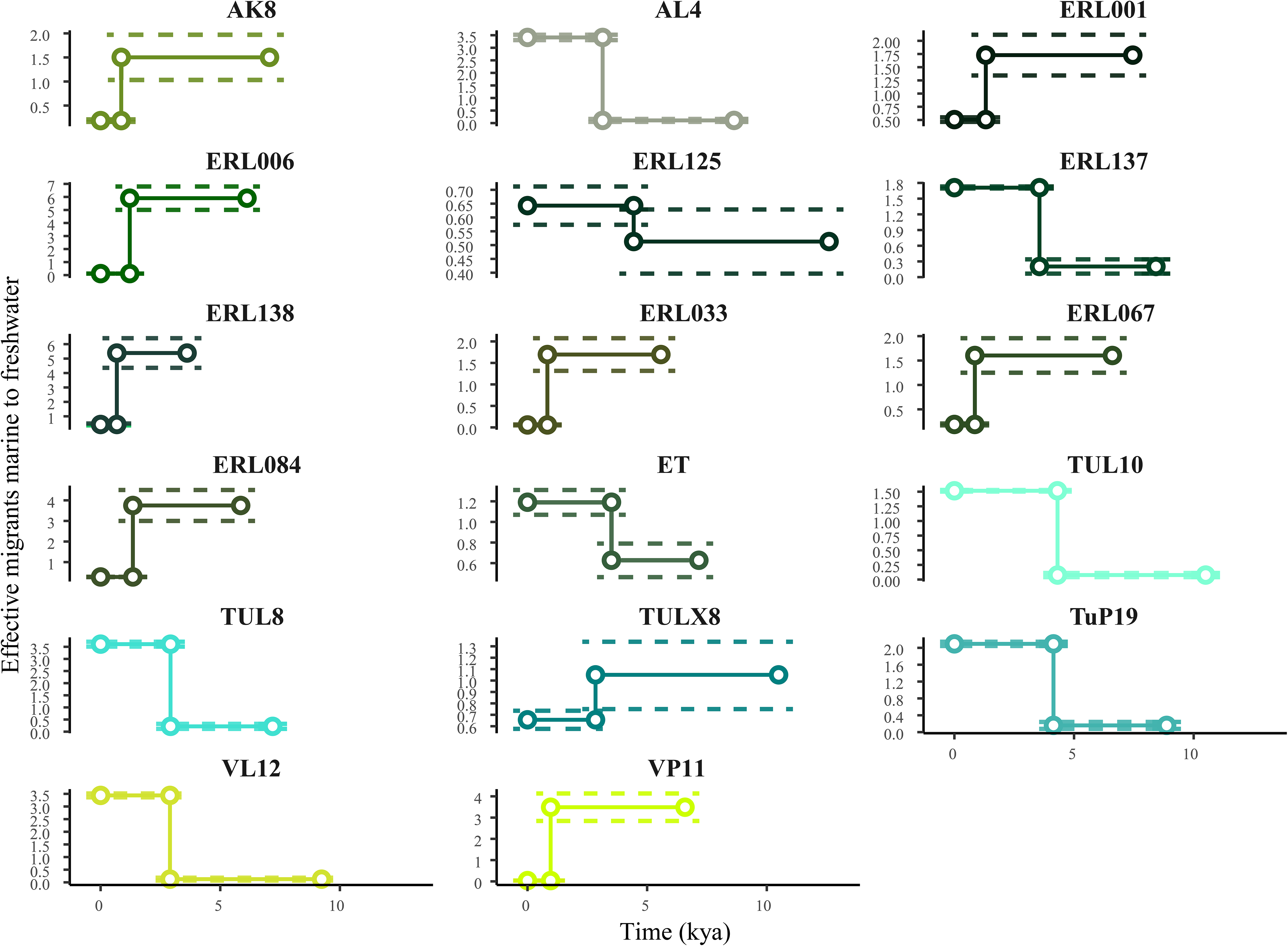
Variation in effective migrants from marine to freshwater for each population. Effective migrants from marine to freshwater are shown against years before present. The first dot for each population on the right represents the time of divergence of the freshwater from the marine; points connected through vertical lines represent the timing of migration change. Dotted lines represent the 95% confidence interval. Geographic locations and population abbreviations in Table S1.

**Figure S8.**
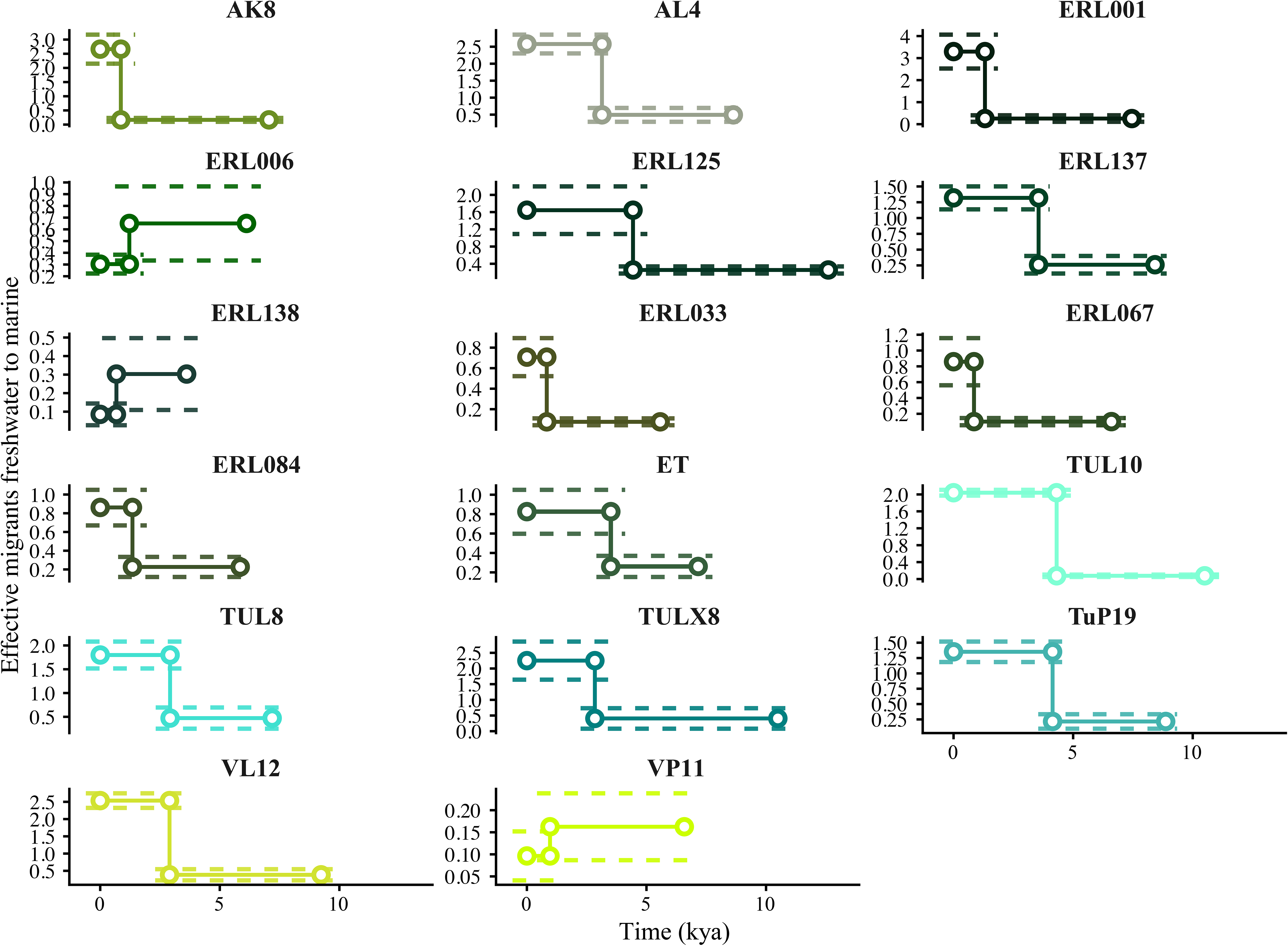
Variation in effective migrants from freshwater to marine for each population. Effective migrants from freshwater to marine are plotted against years before present. The first dot for each population on the right represents the time of divergence of the freshwater from the marine; points connected through vertical lines represent the timing of migration change. Dotted lines represent the 95% confidence interval. Geographic locations and population abbreviations in Table S1.

**Figure S9.**
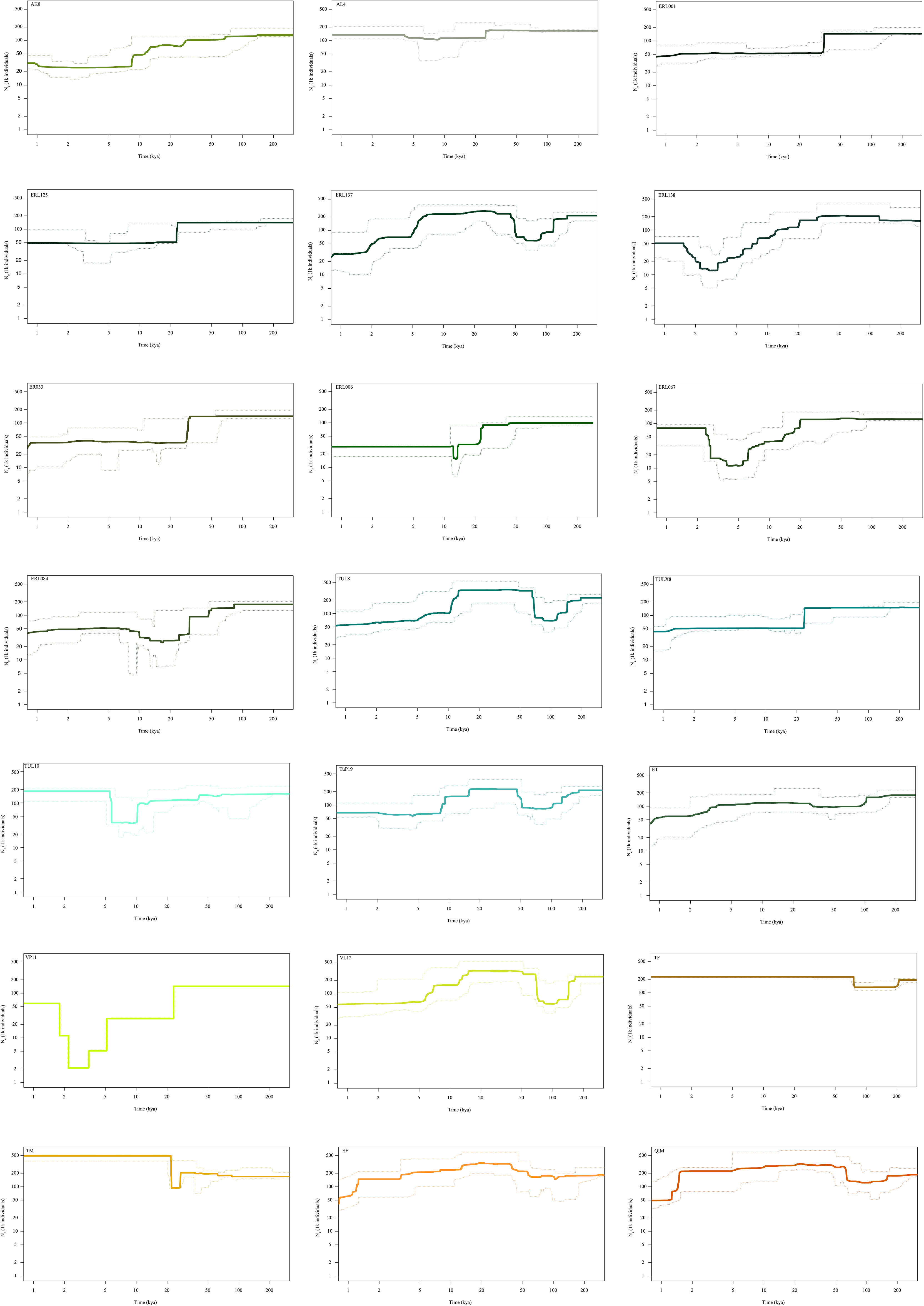
Stairway plots for each population, bold line represents mean and the dotted line the 95% confidence intervals. This is analogous to Figure S6, but with a mutation rate of 5.11×10^−9^. Geographic locations and population abbreviations in Table S1.

**Figure S10.**
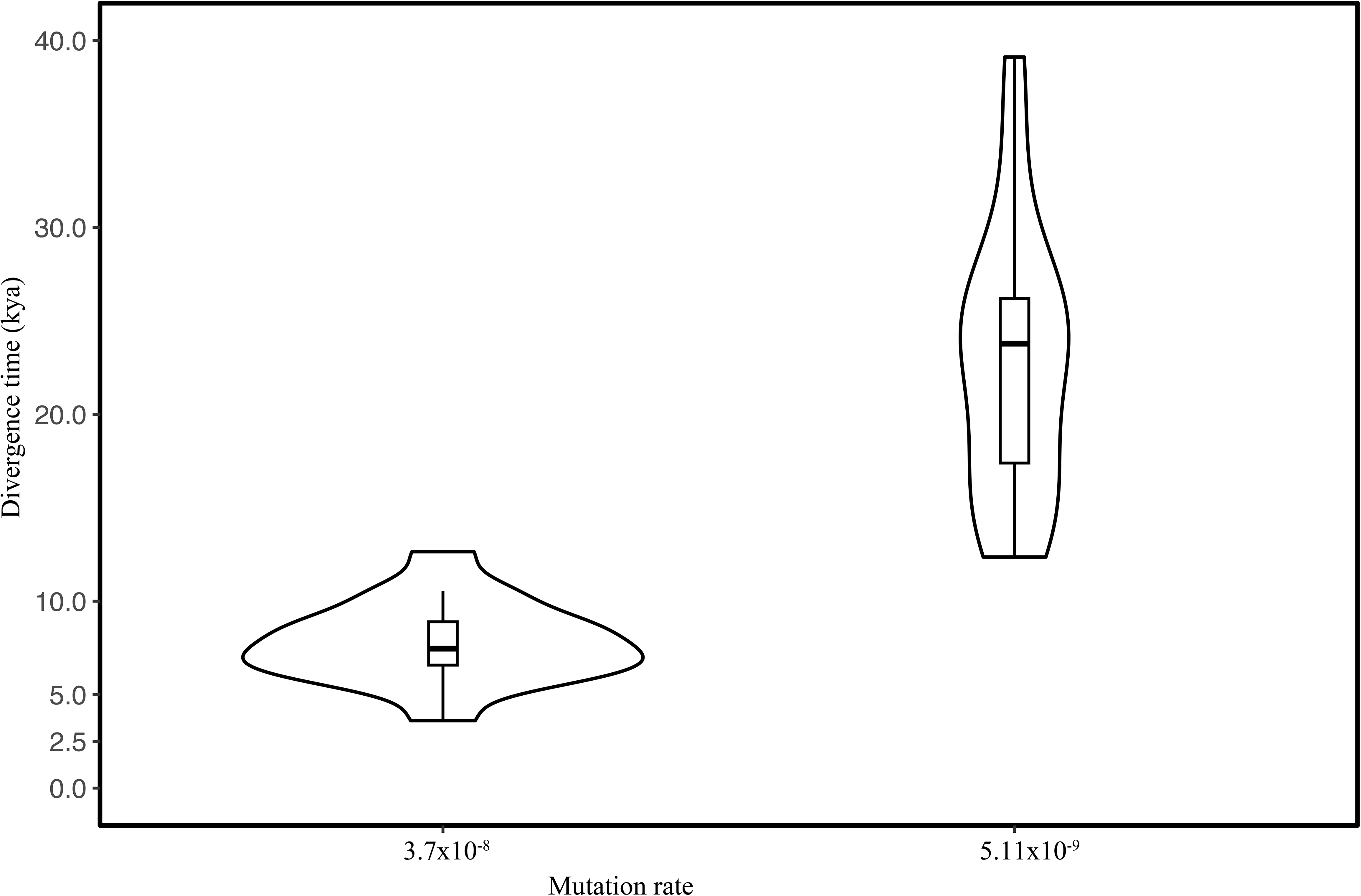
Violin plots of divergence time comparing two different mutation rates. Results of the best run of *fastsimcoal2* modelling are plotted. Mutation rates: 3.7×10^−8^ (used for this study, as well as Liu et al., 2016) and 5.11×10^−9^ (Zhang et al., 2025). Geographic locations and population abbreviations in Table S1.

## Data Accessibility

Raw fastq files of all samples will become accessible under the PRJNA1240318 on NCBI upon publication. Related metadata, scripts, and genotype files will become available as a data package on ERIC, the open and fair Eawag Research Data Institutional Collection (DOI: https://doi.org/10.25678/000E7B) upon publication. All data originates from genetic resources sampled in Greenland under licences G16-040, and G19-002, and any use of the data requires acknowledging Greenland as the country of origin and remains subject to applicable Greenlandic legislation.

## Supporting Information for

**Table S1.** Metadata for sampling location. Location ID, freshwater or marine location, nInd= number of Individuals sampled, Lat=Latitude, Long= Longitude, Altitude, Fish community present in the freshwater Lakes; SB= Sticklebacks, SBC= Sticklebacks and Charr.

| Location ID | Freshwater/marine | nInd | Lat | Long | Altitude | Fish community | Lake Area [ha] | Perimeter [m] | Maximum Depth[m] | Conductivity[μS/cm] |
| --- | --- | --- | --- | --- | --- | --- | --- | --- | --- | --- |
| AK8 | freshwater | 6 | 60.667246 | -46.093646 | 25 | SB | 3.6 | 1287 | 2 | 51 |
| AL4 | freshwater | 6 | 60.668543 | -46.111432 | 14 | SBC | 42.8 | 6090 | 9 | 56 |
| ERL001 | freshwater | 6 | 61.071582 | -45.604472 | 139 | SBC | 51.8 | 10119 | 8 | 65 |
| ERL006 | freshwater | 6 | 61.146044 | -45.528816 | 93 | SB | 7.2 | 1328 | 13 | 146 |
| ERL125 | freshwater | 6 | 61.156861 | -45.636401 | 52 | SB | 3.7 | 1589 | 1.5 | 90 |
| ERL137 | freshwater | 6 | 61.085355 | -45.748242 | 6 | SBC | 13.4 | 1917 | 12 | 87 |
| ERL138 | freshwater | 6 | 61.084701 | -45.732472 | 18 | SB | 6.02 | 1418 | 1 | 77 |
| ERL033 | freshwater | 6 | 61.118369 | -45.580845 | 120 | SBC | 2.5 | 799 | 7 | 110 |
| ERL067 | freshwater | 6 | 61.090048 | -45.653901 | 99 | SBC | 3.2 | 833 | 6 | 70 |
| ERL084 | freshwater | 6 | 61.138109 | -45.586845 | 149 | SB | 2.5 | 611 | 17 | 140 |
| ET | freshwater | 6 | 61.253333 | -45.529141 | 124 | SBC | 26.3 | 4101 | 18 | 34 |
| TUL10 | freshwater | 6 | 60.851511 | -46.463482 | 50 | SB | 64.5 | 5606 | 14 | 25 |
| TUL8 | freshwater | 6 | 60.850441 | -46.493788 | 13 | SBC | 48.1 | 5448 | 20 | 31 |
| TULX8 | freshwater | 6 | 60.862552 | -46.482229 | 36 | SB | 7.3 | 1567 | 5 | 34 |
| TuP19 | freshwater | 6 | 60.824612 | -46.502201 | 4 | SBC | 28.6 | 3542 | 16 | 29 |
| VL12 | freshwater | 6 | 60.724609 | -45.387847 | 26 | SBC | 81.7 | 5059 | na | na |
| VP11 | freshwater | 6 | 60.834834 | -45.334972 | 236 | SB | 8.6 | 1728 | na | na |
| QIM | marine | 6 | 60.709170 | -45.374379 | 0 | na | na | na | na | na |
| Sermilik Fjord (SF) | marine | 7 | 61.116264 | -45.636071 | 0 | na | na | na | na | na |
| Tunulliarfik Fjord (TF) | marine | 3 | 61.243238 | -45.519833 | 0 | na | na | na | na | na |
| Tuttutooq Marine (TM) | marine | 5 | 60.819377 | -46.460235 | 0 | na | na | na | na | na |

**Table S2.** Individual sample IDs, their population affiliation, sequencing depth, sequencing depth standard deviation and NCBI accession number are presented. Geographic locations and population abbreviations can be found in Table S1.

| Individual ID | Population | Depth | Depth sd | NCBI Accession number |
| --- | --- | --- | --- | --- |
| 101838 | VP11 | 17.70 | 4.57 | SAMN47516354 |
| 101839 | VP11 | 22.18 | 5.23 | SAMN47516355 |
| 101840 | VP11 | 23.83 | 5.38 | SAMN47516356 |
| 101841 | VP11 | 14.49 | 4.35 | SAMN47516357 |
| 101842 | VP11 | 33.99 | 6.48 | SAMN47516358 |
| 101843 | VP11 | 23.44 | 5.29 | SAMN47516359 |
| 141591 | QIM | 20.16 | 4.82 | SAMN47516360 |
| 141592 | QIM | 19.58 | 4.71 | SAMN47516361 |
| 141594 | QIM | 21.08 | 5.07 | SAMN47516362 |
| 141595 | QIM | 19.79 | 4.78 | SAMN47516363 |
| 141596 | QIM | 19.93 | 4.79 | SAMN47516364 |
| 141597 | QIM | 23.78 | 5.31 | SAMN47516365 |
| 142105 | VL12 | 18.49 | 4.69 | SAMN47516366 |
| 142106 | VL12 | 20.16 | 4.94 | SAMN47516367 |
| 142107 | VL12 | 13.98 | 4.28 | SAMN47516368 |
| 142108 | VL12 | 20.84 | 5.04 | SAMN47516369 |
| 142109 | VL12 | 31.69 | 6.30 | SAMN47516370 |
| 142110 | VL12 | 15.13 | 4.18 | SAMN47516371 |
| 185157 | ET | 9.97 | 3.27 | SAMN47516372 |
| 185167 | ET | 11.03 | 3.45 | SAMN47516373 |
| 185175 | ET | 10.49 | 3.36 | SAMN47516374 |
| 185182 | ET | 9.97 | 3.27 | SAMN47516375 |
| 185187 | ET | 9.75 | 3.24 | SAMN47516376 |
| 185191 | ET | 10.17 | 3.32 | SAMN47516377 |
| 185203 | ERL006 | 10.34 | 3.35 | SAMN47516378 |
| 185230 | ERL006 | 11.04 | 3.50 | SAMN47516379 |
| 185236 | ERL006 | 9.90 | 3.32 | SAMN47516380 |
| 185239 | ERL006 | 9.87 | 3.30 | SAMN47516381 |
| 185240 | ERL006 | 10.26 | 3.36 | SAMN47516382 |
| 185243 | ERL006 | 10.16 | 3.34 | SAMN47516383 |
| 185258 | AL4 | 10.68 | 3.43 | SAMN47516384 |
| 185263 | AL4 | 11.10 | 3.46 | SAMN47516385 |
| 185264 | AL4 | 11.37 | 3.51 | SAMN47516386 |
| 185277 | AL4 | 9.33 | 3.16 | SAMN47516387 |
| 185282 | AL4 | 10.78 | 3.41 | SAMN47516388 |
| 185291 | AL4 | 10.08 | 3.29 | SAMN47516389 |
| 185399 | Ak8 | 9.73 | 3.24 | SAMN47516390 |
| 185406 | Ak8 | 12.20 | 3.65 | SAMN47516391 |
| 185411 | Ak8 | 10.31 | 3.33 | SAMN47516392 |
| 185428 | Ak8 | 10.31 | 3.34 | SAMN47516393 |
| 185431 | Ak8 | 10.91 | 3.44 | SAMN47516394 |
| 185432 | Ak8 | 9.51 | 3.18 | SAMN47516395 |
| 185498 | ERL33 | 10.98 | 3.46 | SAMN47516396 |
| 185511 | ERL33 | 10.04 | 3.29 | SAMN47516397 |
| 185512 | ERL33 | 10.83 | 3.43 | SAMN47516398 |
| 185523 | ERL33 | 10.02 | 3.29 | SAMN47516399 |
| 185524 | ERL33 | 10.59 | 3.40 | SAMN47516400 |
| 185525 | ERL33 | 10.72 | 3.41 | SAMN47516401 |
| 185556 | ERL001 | 9.18 | 3.19 | SAMN47516402 |
| 185563 | ERL001 | 9.64 | 3.27 | SAMN47516403 |
| 185571 | ERL001 | 9.35 | 3.21 | SAMN47516404 |
| 185572 | ERL001 | 11.28 | 3.55 | SAMN47516405 |
| 185577 | ERL001 | 9.89 | 3.32 | SAMN47516406 |
| 185584 | ERL001 | 10.97 | 3.50 | SAMN47516407 |
| 186568 | Tunulliarfik Fjord | 25.72 | 5.62 | SAMN47516408 |
| 186570 | Tunulliarfik Fjord | 10.52 | 3.37 | SAMN47516409 |
| 186572 | Tunulliarfik Fjord | 14.30 | 3.99 | SAMN47516410 |
| 186573 | Tunulliarfik Fjord | 11.35 | 3.51 | SAMN47516411 |
| 186575 | Sermilik Fjord | 28.07 | 5.81 | SAMN47516412 |
| 186577 | Sermilik Fjord | 16.93 | 4.38 | SAMN47516413 |
| 186578 | Sermilik Fjord | 10.55 | 3.37 | SAMN47516414 |
| 186579 | Sermilik Fjord | 11.37 | 3.50 | SAMN47516415 |
| 186582 | Sermilik Fjord | 25.73 | 5.57 | SAMN47516416 |
| 186583 | Sermilik Fjord | 19.38 | 4.73 | SAMN47516417 |
| 186751 | ERL67 | 11.67 | 3.58 | SAMN47516418 |
| 186752 | ERL67 | 9.64 | 3.25 | SAMN47516419 |
| 186766 | ERL67 | 9.45 | 3.19 | SAMN47516420 |
| 186771 | ERL67 | 10.17 | 3.32 | SAMN47516421 |
| 186787 | ERL67 | 10.11 | 3.37 | SAMN47516422 |
| 186790 | ERL67 | 9.48 | 3.20 | SAMN47516423 |
| 186848 | Tuttutooq Marine | 17.91 | 4.55 | SAMN47516424 |
| 188104 | TULX8 | 10.38 | 3.33 | SAMN47516425 |
| 188107 | TULX8 | 10.01 | 3.27 | SAMN47516426 |
| 188115 | TULX8 | 10.16 | 3.30 | SAMN47516427 |
| 188120 | TULX8 | 10.93 | 3.44 | SAMN47516428 |
| 188122 | TULX8 | 10.76 | 3.40 | SAMN47516429 |
| 188146 | TULX8 | 11.16 | 3.47 | SAMN47516430 |
| 188968 | Tuttutooq Marine | 36.01 | 6.77 | SAMN47516431 |
| 188969 | Tuttutooq Marine | 22.48 | 5.18 | SAMN47516432 |
| 190009 | ERL84 | 10.09 | 3.31 | SAMN47516433 |
| 190010 | ERL84 | 10.59 | 3.39 | SAMN47516434 |
| 190018 | ERL84 | 10.18 | 3.30 | SAMN47516435 |
| 190024 | ERL84 | 10.06 | 3.30 | SAMN47516436 |
| 190025 | ERL84 | 10.37 | 3.35 | SAMN47516437 |
| 190041 | ERL84 | 11.32 | 3.52 | SAMN47516438 |
| 190202 | TuP19 | 10.85 | 3.42 | SAMN47516439 |
| 190214 | TuP19 | 10.18 | 3.32 | SAMN47516440 |
| 190224 | TuP19 | 10.93 | 3.43 | SAMN47516441 |
| 190226 | TuP19 | 9.82 | 3.24 | SAMN47516442 |
| 190238 | TuP19 | 11.09 | 3.48 | SAMN47516443 |
| 190248 | TuP19 | 11.86 | 3.58 | SAMN47516444 |
| 190258 | ERL137 | 9.67 | 3.26 | SAMN47516445 |
| 190260 | ERL137 | 9.97 | 3.31 | SAMN47516446 |
| 190280 | ERL137 | 11.02 | 3.50 | SAMN47516447 |
| 190289 | ERL137 | 10.47 | 3.42 | SAMN47516448 |
| 190291 | ERL137 | 10.32 | 3.37 | SAMN47516449 |
| 190293 | ERL137 | 10.03 | 3.32 | SAMN47516450 |
| 190406 | ERL138 | 9.39 | 3.17 | SAMN47516451 |
| 190409 | ERL138 | 10.65 | 3.38 | SAMN47516452 |
| 190420 | ERL138 | 10.53 | 3.39 | SAMN47516453 |
| 190421 | ERL138 | 10.41 | 3.36 | SAMN47516454 |
| 190440 | ERL138 | 11.44 | 3.53 | SAMN47516455 |
| 190445 | ERL138 | 10.22 | 3.34 | SAMN47516456 |
| 190655 | ERL125 | 10.04 | 3.29 | SAMN47516457 |
| 190667 | ERL125 | 10.88 | 3.45 | SAMN47516458 |
| 190669 | ERL125 | 11.50 | 3.57 | SAMN47516459 |
| 190673 | ERL125 | 13.10 | 3.85 | SAMN47516460 |
| 190692 | ERL125 | 8.92 | 3.14 | SAMN47516461 |
| 190696 | ERL125 | 9.56 | 3.22 | SAMN47516462 |
| 190723 | TUL10 | 10.09 | 3.30 | SAMN47516463 |
| 190727 | TUL10 | 10.58 | 3.37 | SAMN47516464 |
| 190728 | TUL10 | 10.21 | 3.32 | SAMN47516465 |
| 190733 | TUL10 | 10.53 | 3.37 | SAMN47516466 |
| 190739 | TUL10 | 10.66 | 3.41 | SAMN47516467 |
| 190740 | TUL10 | 10.60 | 3.38 | SAMN47516468 |
| 190813 | TUL8 | 12.60 | 3.72 | SAMN47516469 |
| 190814 | TUL8 | 10.55 | 3.40 | SAMN47516470 |
| 190823 | TUL8 | 9.99 | 3.29 | SAMN47516471 |
| 190831 | TUL8 | 10.97 | 3.44 | SAMN47516472 |
| 190833 | TUL8 | 11.02 | 3.45 | SAMN47516473 |
| 190844 | TUL8 | 9.71 | 3.24 | SAMN47516474 |
| 204160 | Tuttutooq Marine | 10.38 | 3.39 | SAMN47516475 |
| 204161 | Tuttutooq Marine | 11.35 | 3.54 | SAMN47516476 |
| 219604 | SF | 22.82 | 5.20 | SAMN47516477 |

**Table S3.** Overview of filtering steps and reduction of SNP and invariants sites at each step as well as total number of sites.

| Filter | SNP | Invariant sites | Total number of sites |
| --- | --- | --- | --- |
| No filter | 8030216 | 336063657 | 344093873 |
| QUAL 20 | 7744436 | 193404643 | 201149079 |
| Coverage 4 x | 7343661 | 186584539 | 193928200 |
| mean depth | 5984917 | 161151909 | 167136826 |
| Max miss 0.5 | 5984760 | 161150045 | 167134805 |
| Biallelic SNPs | 5928003 | 161150045 | 167078048 |
| 10 bp SNP thinned | 4723738 | 161150045 | 165873783 |

**Table S4.**
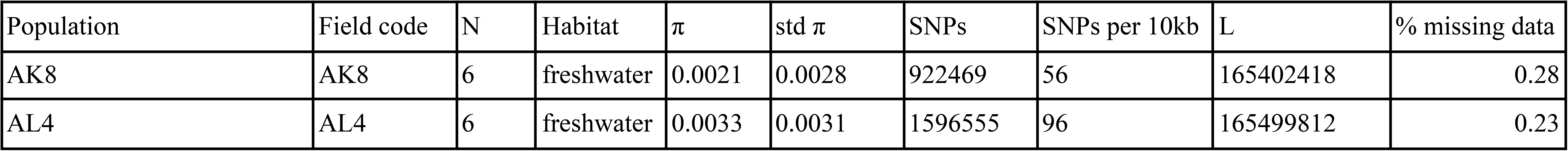
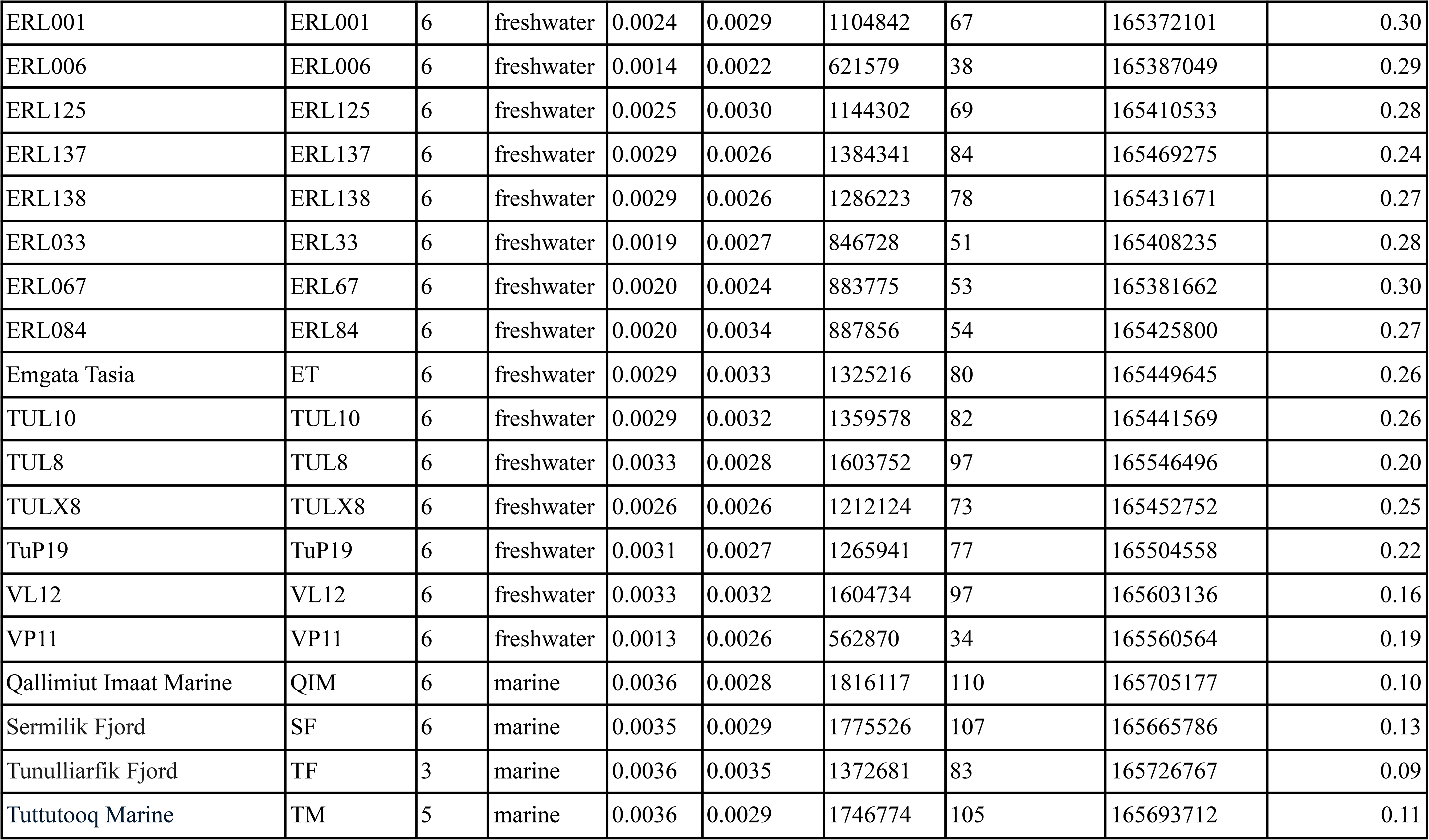
Population statistics of sampling locations. The number of individuals (N) used for calculation of the statistic, type of habitat, nucleotide diversity (π) and its standard variation as well as total number of SNPs and SNPs per 10kb, the total number of sites for 1d sfs calculation (L) and the percent of missing data per population with a maximum missing filter of 1 (note Table S3 number of total sites is the base for this calculation). Geographic locations and population abbreviations can be found in Table S1.

| Population | Field code | N | Habitat | $\pi$ | std $\pi$ | SNPs | SNPs per 10kb | L | % missing data |
| --- | --- | --- | --- | --- | --- | --- | --- | --- | --- |
| AK8 | AK8 | 6 | freshwater | 0.0021 | 0.0028 | 922469 | 56 | 165402418 | 0.28 |
| AL4 | AL4 | 6 | freshwater | 0.0033 | 0.0031 | 1596555 | 96 | 165499812 | 0.23 |
| ERL001 | ERL001 | 6 | freshwater | 0.0024 | 0.0029 | 1104842 | 67 | 165372101 | 0.30 |
| ERL006 | ERL006 | 6 | freshwater | 0.0014 | 0.0022 | 621579 | 38 | 165387049 | 0.29 |
| ERL125 | ERL125 | 6 | freshwater | 0.0025 | 0.0030 | 1144302 | 69 | 165410533 | 0.28 |
| ERL137 | ERL137 | 6 | freshwater | 0.0029 | 0.0026 | 1384341 | 84 | 165469275 | 0.24 |
| ERL138 | ERL138 | 6 | freshwater | 0.0029 | 0.0026 | 1286223 | 78 | 165431671 | 0.27 |
| ERL033 | ERL33 | 6 | freshwater | 0.0019 | 0.0027 | 846728 | 51 | 165408235 | 0.28 |
| ERL067 | ERL67 | 6 | freshwater | 0.0020 | 0.0024 | 883775 | 53 | 165381662 | 0.30 |
| ERL084 | ERL84 | 6 | freshwater | 0.0020 | 0.0034 | 887856 | 54 | 165425800 | 0.27 |
| Emgata Tasia | ET | 6 | freshwater | 0.0029 | 0.0033 | 1325216 | 80 | 165449645 | 0.26 |
| TUL10 | TUL10 | 6 | freshwater | 0.0029 | 0.0032 | 1359578 | 82 | 165441569 | 0.26 |
| TUL8 | TUL8 | 6 | freshwater | 0.0033 | 0.0028 | 1603752 | 97 | 165546496 | 0.20 |
| TULX8 | TULX8 | 6 | freshwater | 0.0026 | 0.0026 | 1212124 | 73 | 165452752 | 0.25 |
| TuP19 | TuP19 | 6 | freshwater | 0.0031 | 0.0027 | 1265941 | 77 | 165504558 | 0.22 |
| VL12 | VL12 | 6 | freshwater | 0.0033 | 0.0032 | 1604734 | 97 | 165603136 | 0.16 |
| VP11 | VP11 | 6 | freshwater | 0.0013 | 0.0026 | 562870 | 34 | 165560564 | 0.19 |
| Qallimiut Imaat Marine | QIM | 6 | marine | 0.0036 | 0.0028 | 1816117 | 110 | 165705177 | 0.10 |
| Sermilik Fjord | SF | 6 | marine | 0.0035 | 0.0029 | 1775526 | 107 | 165665786 | 0.13 |
| Tunulliarfik Fjord | TF | 3 | marine | 0.0036 | 0.0035 | 1372681 | 83 | 165726767 | 0.09 |
| Tuttutooq Marine | TM | 5 | marine | 0.0036 | 0.0029 | 1746774 | 105 | 165693712 | 0.11 |

**Table S5.**
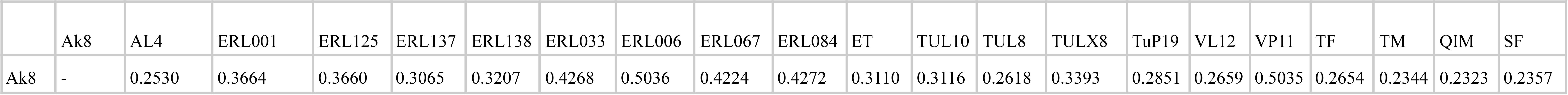
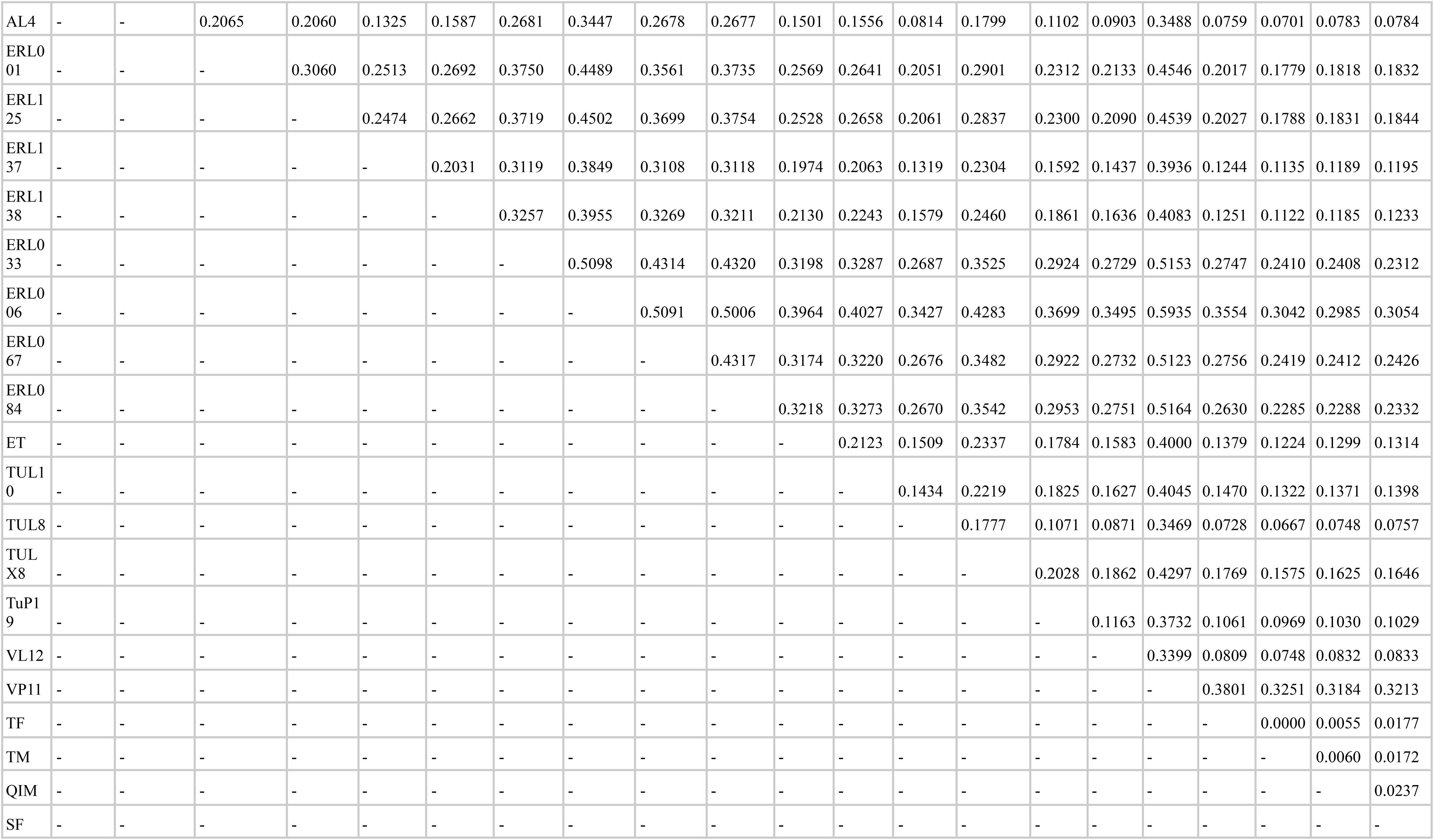
Pairwise F_ST_ of sampling locations. Geographic locations and population abbreviations can be found in Table S1.

**Table S6.** Demographic parameter estimates vary among populations. Results of the bootstrapping of the final model (model g) of *fastsimcoal2* with a mutation rate of 3.7× 10^−8^ and K= 9, which is the number of parameters in the *fastsimcoal2* model. All parameters are mean values of the bootstrapping. Effective population sizes (N_e_) of ancestral, freshwater, marine, N_1_M_21rec_= effective migrants from marine to freshwater after CHANGEM (present), N_1_M_21old_= effective migrants from marine to freshwater before CHANGEM (past), N_2_M_12rec_= effective migrants from freshwater to marine after CHANGEM (present), N_2_M_12old_= effective migrants from freshwater to marine before CHANGEM (past), T_DIV_= divergence time in kya, CHANGEM= timing of migration matrix change in kya, MIG_21_= migration from marine to freshwater, MIG_12_= migration from freshwater to marine; MaxEstLhood= maximum estimated likelihood, MaxObsLhood= maximum observed likelihood, AIC= Akaike information criterion, delta= difference of MaxEstLhood and MaxObsLhood. Geographic locations and population abbreviations can be found in Table S1.

| Population | $N_e$ ancestral | $N_e$ freshwater | $N_e$ marine | $N_1M_{21rec}$ | $N_2M_{12rec}$ | $N_1M_{21old}$ | $N_2M_{12old}$ | $T_{DIV}$ | CHANGM | $MIG_{21rec} \times 10^{-04}$ | $MIG_{12rec} \times 10^{-04}$ | $MIG_{21old} \times 10^{-04}$ | $MIG_{12old} \times 10^{-04}$ | MaxEstLhood | MaxObsLhood | K | AIC | delta |
| --- | --- | --- | --- | --- | --- | --- | --- | --- | --- | --- | --- | --- | --- | --- | --- | --- | --- | --- |
| AK8 | 47519 | 2676 | 56165 | 0.19 | 2.66 | 1.50 | 0.17 | 7.06 | 0.86 | 0.70 | 0.48 | 5.60 | 0.03 | -7846512 | -7845178 | 9 | 36134541 | -1334 |
| AL4 | 47625 | 11109 | 49009 | 3.41 | 2.58 | 0.11 | 0.49 | 8.63 | 3.14 | 3.20 | 0.54 | 0.10 | 0.10 | -8723930 | -8708901 | 9 | 40175199 | -15029 |
| ERL001 | 47398 | 3690 | 56314 | 0.51 | 3.30 | 1.73 | 0.26 | 7.46 | 1.31 | 1.37 | 0.58 | 4.68 | 0.005 | -8060737 | -8059213 | 9 | 37121083 | -1524 |
| ERL006 | 47133 | 1192 | 64188 | 0.12 | 0.30 | 5.91 | 0.65 | 6.12 | 1.22 | 0.98 | 0.50 | 49.08 | 0.01 | -7343219 | -7342548 | 9 | 33816789 | -670 |
| ERL125 | 46877 | 3272 | 52355 | 0.65 | 1.64 | 0.51 | 0.25 | 12.65 | 4.42 | 2.01 | 0.31 | 1.52 | 0.05 | -8156239 | -8153465 | 9 | 37560888 | -2774 |
| ERL137 | 47489 | 5311 | 53385 | 1.71 | 1.32 | 0.20 | 0.26 | 8.42 | 3.56 | 0.33 | 0.25 | 0.37 | 0.05 | -8451638 | -8443669 | 9 | 38921251 | -7970 |
| ERL138 | 48112 | 3680 | 64089 | 0.44 | 0.09 | 5.39 | 0.30 | 3.61 | 0.68 | 1.31 | 0.14 | 14.98 | 0.05 | -8196876 | -8194823 | 9 | 37748027 | -2053 |
| ERL033 | 47929 | 2300 | 57618 | 0.06 | 0.71 | 1.69 | 0.08 | 5.57 | 0.83 | 0.25 | 0.13 | 7.20 | 0.01 | -7740624 | -7739264 | 9 | 35646910 | -1360 |
| ERL067 | 47650 | 2110 | 57710 | 0.19 | 0.86 | 1.60 | 0.10 | 6.60 | 0.85 | 0.90 | 0.16 | 7.53 | 0.02 | -7774087 | -7772895 | 9 | 35801013 | -1192 |
| ERL084 | 47411 | 2126 | 62473 | 0.28 | 0.86 | 3.75 | 0.23 | 5.85 | 1.35 | 1.32 | 0.15 | 0.002 | 0.04 | -7731983 | -7731151 | 9 | 35607117 | -832 |
| ET | 47614 | 4629 | 54761 | 1.18 | 1.53 | 0.69 | 0.44 | 7.92 | 3.56 | 2.57 | 0.28 | 1.49 | 0.08 | -8343989 | -8339568 | 9 | 38425505 | -4420 |
| TUL10 | 46962 | 6756 | 49530 | 1.52 | 2.04 | 0.08 | 0.08 | 10.53 | 4.31 | 2.25 | 0.41 | 0.12 | 0.02 | -8434637 | -8429042 | 9 | 38842957 | -5595 |
| TUL8 | 47647 | 10147 | 51835 | 3.61 | 1.80 | 0.22 | 0.47 | 7.19 | 2.92 | 3.68 | 0.37 | 0.21 | 0.09 | -8707994 | -8698846 | 9 | 40101812 | -9148 |
| TULX8 | 47582 | 3777 | 53276 | 0.66 | 2.25 | 1.05 | 0.41 | 10.49 | 2.84 | 1.76 | 0.42 | 2.66 | 0.08 | -8231055 | -8228243 | 9 | 37905428 | -2812 |
| TuP19 | 47714 | 6674 | 51086 | 2.09 | 1.36 | 0.16 | 0.21 | 8.90 | 4.14 | 3.22 | 0.27 | 0.24 | 0.04 | -8561344 | -8549851 | 9 | 39426465 | -11493 |
| VL12 | 47575 | 11372 | 48527 | 3.44 | 2.54 | 0.13 | 0.38 | 9.24 | 2.90 | 3.09 | 0.53 | 0.11 | 0.08 | -8767756 | -8756223 | 9 | 40377028 | -11533 |
| VP11 | 47673 | 1084 | 60685 | 0.04 | 0.10 | 3.49 | 0.16 | 6.58 | 0.97 | 0.34 | 0.17 | 30.40 | 0.03 | -7304472 | -7303078 | 9 | 33638354 | -1394 |

**Table S7.** Fastsimcoal2 output for models a-g with a mutation rate of 3.7 x 10^−8^. Effective population sizes (N_e_) of ancestral, freshwater, marine,TDIV= time of divergence, CHANGEM= timing of migration matrix change, MIG21= migration from marine to freshwater, MIG12= migration from freshwater to marine; MaxEstLhood= maximum estimated likelihood, MaxObsLhood= maximum observed likelihood, K= number of parameters in fastSimCoal modelling, AIC= Akaike information criterion, delta= difference of MaxEstLhood and MaxObsLhood, NBOT= number of individuals at population decline, TBOT= timing of population decline, NEXP= number of individuals at population expansion, TEXP= timing of population expansion, N1M21rec= number of individuals migrating from marine to freshwater after CHANGEM (present), N1M21old= number of individuals migrating from marine to freshwater before CHANGEM (past), N2M12rec= number of individuals migrating from freshwater to marine after CHANGEM (present), N2M12old= number of individuals migrating from freshwater to marine before CHANGEM (past). Geographic locations and population abbreviations can be found in Table S1. Table S7 is supplied as a separate Table_S7.csv file.

**Table S8.** Demographic parameter estimates vary among populations. Results of the bootstrapping of the final model (model g) of *fastsimcoal2* with a mutation rate of 5.11×10^−9^ and K= 9, which is the number of parameters in the *fastsimcoal2* model. All parameters are mean values of the bootstrapping. Effective population sizes (N_e_) of ancestral, freshwater, marine, N_1_M_21rec_= effective migrants from marine to freshwater after CHANGEM (present), N_1_M_21old_= effective migrants from marine to freshwater before CHANGEM (past), N_2_M_12rec_= effective migrants from freshwater to marine after CHANGEM (present), N_2_M_12old_= effective migrants from freshwater to marine before CHANGEM (past), T_DIV_= divergence time in kya, CHANGEM= timing of migration matrix change in kya, MIG_21_= migration from marine to freshwater, MIG_12_= migration from freshwater to marine; MaxEstLhood= maximum estimated likelihood, MaxObsLhood= maximum observed likelihood, AIC= Akaike information criterion, delta= difference of MaxEstLhood and MaxObsLhood. Geographic locations and population abbreviations can be found in Table S1. Table S8 is supplied as a separate Table_S8.csv file.

**Table S9.** Demographic parameter estimates vary among populations and for different outgroups. Results of the bootstrapping of the final model (model g) of *fastsimcoal2* with a mutation rate of 3.7 x 10^−8^ and K= 9, which is the number of parameters in the *fastsimcoal2* model. All parameters are mean values of the bootstrapping. Effective population sizes (N_e_) of ancestral, freshwater, marine, N_1_M_21rec_= effective migrants from marine to freshwater after CHANGEM (present), N_1_M_21old_= effective migrants from marine to freshwater before CHANGEM (past), N_2_M_12rec_= effective migrants from freshwater to marine after CHANGEM (present), N_2_M_12old_= effective migrants from freshwater to marine before CHANGEM (past), T_DIV_= divergence time in kya, CHANGEM= timing of migration matrix change in kya, MIG_21_= migration from marine to freshwater, MIG_12_= migration from freshwater to marine; MaxEstLhood= maximum estimated likelihood, MaxObsLhood= maximum observed likelihood, AIC= Akaike information criterion, delta= difference of MaxEstLhood and MaxObsLhood. Geographic locations and population abbreviations can be found in Table S1.

| Population | $N_e$ ancestral | $N_e$ freshwater | $N_e$ marine | $N_1M_{21rec}$ | $N_2M_{12rec}$ | $N_1M_{21old}$ | $N_2M_{12old}$ | $T_{DIV}$ | CHANGM | $MIG_{21rec} \times 10^{-04}$ | $MIG_{12rec} \times 10^{-04}$ | $MIG_{21old} \times 10^{-04}$ | $MIG_{12old} \times 10^{-04}$ | MaxEstLhood | MaxObsLhood |
| --- | --- | --- | --- | --- | --- | --- | --- | --- | --- | --- | --- | --- | --- | --- | --- |
| AL4 & mixed marine | 47725 | 10555 | 57381 | 3.52 | 2.67 | 0.14 | 0.70 | 6569 | 2702 | 3.55 | 0.51 | 0.13 | 0.13 | -8729040 | -8716676 |
| AL4 & QIM | 47625 | 11109 | 49009 | 3.41 | 2.58 | 0.11 | 0.49 | 8631 | 3140 | 3.20 | 0.54 | 0.10 | 0.10 | -8723930 | -8708901 |
| ERL138 & mixed marine | 49942 | 5443 | 65048 | 1.23 | 3.75 | 1.61 | 2.67 | 9183 | 2464 | 11.67 | 0.73 | 3.78 | 0.46 | -1481109 | -1440887 |
| ERL138 & QIM | 48112 | 3680 | 64089 | 0.44 | 0.09 | 5.39 | 0.30 | 3610 | 0.68 | 1.31 | 0.14 | 14.98 | 0.05 | -8196876 | -8194823 |
| ET & mixed marine | 47984 | 4262 | 67555 | 1.14 | 1.47 | 0.83 | 0.80 | 5028 | 2385 | 2.71 | 0.23 | 1.99 | 0.13 | -8352409 | -8349122 |
| ET & QIM | 47614 | 4629 | 54761 | 1.18 | 1.53 | 0.69 | 0.44 | 7924 | 3560 | 2.57 | 0.28 | 1.49 | 0.08 | -8343989 | -8339568 |
| TUL10 & mixed marine | 46630 | 6433 | 54129 | 1.55 | 2.49 | 0.10 | 0.09 | 10048 | 3859 | 2.43 | 0.46 | 0.16 | 0.02 | -8456051 | -8451679 |
| TUL10 & QIM | 46962 | 6756 | 49530 | 1.52 | 2.04 | 0.08 | 0.08 | 10530 | 4306 | 2.25 | 0.41 | 0.12 | 0.02 | -8434637 | -8429042 |
| VL12 & mixed marine | 47298 | 10609 | 54895 | 3.69 | 3.10 | 0.09 | 0.40 | 7721 | 2502 | 3.58 | 0.58 | 0.09 | 0.07 | -8766978 | -8757754 |
| VL12 & QIM | 47575 | 11372 | 48527 | 3.44 | 2.54 | 0.13 | 0.38 | 9235 | 2898 | 3.09 | 0.53 | 0.11 | 0.08 | -8767756 | -8756223 |

